# Reefal regions were biodiversity hotspots throughout the Phanerozoic

**DOI:** 10.1101/2025.01.16.633324

**Authors:** Roger A. Close, Roger B.J. Benson, Wolfgang Kiessling, Erin E. Saupe

## Abstract

Reefs are important hotspots of marine biodiversity today, and acted as cradles of diversification in the geological past. However, we know little about how the diversity of reef-supporting regions varied through deep time, and how this differed from other regions. We quantified regional diversity patterns in reef-supporting and non-reef-supporting regions in the fossil record of Phanerozoic marine invertebrates. Diversity in reef-supporting regions is on average two- to three-fold higher than in non-reef-supporting regions, and has been remarkably stable over timescales of tens to hundreds of millions of years. This signal is present in both reefal and non-reefal facies within reef-supporting regions, suggesting that reefs enriched diversity in surrounding environments. Sepkoski’s ‘Modern Fauna’, an assemblage of higher taxa that includes gastropods, bivalves and echinoids, has been a key component of reef-supporting regions since the Paleozoic, contrasting with its later rise to dominance in non-reef-supporting regions during the later Mesozoic–Cenozoic.

**One-Sentence Summary:** Regions of the globe that supported reefal environments have been key hotspots of marine animal diversity for over 400 million years.

## Main Text

Biodiversity varies considerably among environments on Earth today. In the oceans, tropical coastal zones are approximately twice as rich as temperate coastal zones (*1, 2*), and coral-algal reefs host a disproportionate share of extant biodiversity, supporting an estimated one-third of all marine species despite occupying only 0.1% of Earth’s surface (*1, 3*). Diversity also varies among environments in the fossil record [e.g., refs (*4, 5*)]. Regionally, tropical shallow-water habitats, especially reefs, have been sites of net origination, exporting species to other environments and to higher latitudes in deep time (*6,7)*). Indeed, the occurrence of high diversity within reefal environments is a persistent, first-order observation of paleobiology [e.g., refs (*8–13*)]. Geographic regions that support reefal environments may therefore be characterised by different diversity dynamics to those that do not support reefs on Phanerozoic timescales. These differences may be central to explaining large-scale patterns in the taxonomic composition and richness of the marine biota [e.g., refs (*14, 15*)], but they remain poorly understood, and no study has directly contrasted diversity within reefal and non-reefal environments over the Phanerozoic.

Patterns of marine animal diversity through the Phanerozoic have long been analyzed at a notionally ‘global’ scale. However, the fossil record for any time in Earth’s history is never truly global in scope, with sampled geographic regions varying substantially in number, size, location, and environmental context. Therefore, much of the apparent variation in global diversity through time may be structured by variation in the geographic and environmental scope of the sampled fossil record (*5, 8, 10, 16, 17*). Spatially- and environmentally-explicit analyses can overcome these biases, and have great potential for uncovering variation in the geographic and environmental structure of diversity through time. Here, we quantify changes in diversity for reef-supporting versus non-reef-supporting regions over the Phanerozoic, to identify hotspots of biodiversity on multi-million-year timescales and assess spatial variation in macroevolutionary patterns.

### Fossil occurrence data and reconstruction of diversity among environments

We reconstructed patterns of local- to regional-scale diversity at genus level for Phanerozoic marine animals using occurrence data from the Paleobiology Database (*18, 19*). We present results for a range of sifting criteria, but focus on those that exclude deposits that are unlithified or poorly-lithified and sieved, or which lack information about lithification style (*19*). To estimate diversity at smaller spatiotemporal scales, we tallied counts of genera per collection (i.e., spatially- and temporally-resolved fossil localities; a measure of alpha diversity, or within-community/local richness), and per geological formation (*19*). To obtain regional-scale diversity estimates while standardizing for spatial sampling, we rasterized fossil localities into equal-area hexagonal/pentagonal grid cells with 100 km, 500 km, 1000 km, and 2000 km spacings [our main-text results focus on 1000 km spacings; Fig. 1 and fig. S1; (*19*)], before computing diversity for each grid cell using Shareholder Quorum Subsampling [SQS (*15, 19, 20*)].

**Fig. 1:**
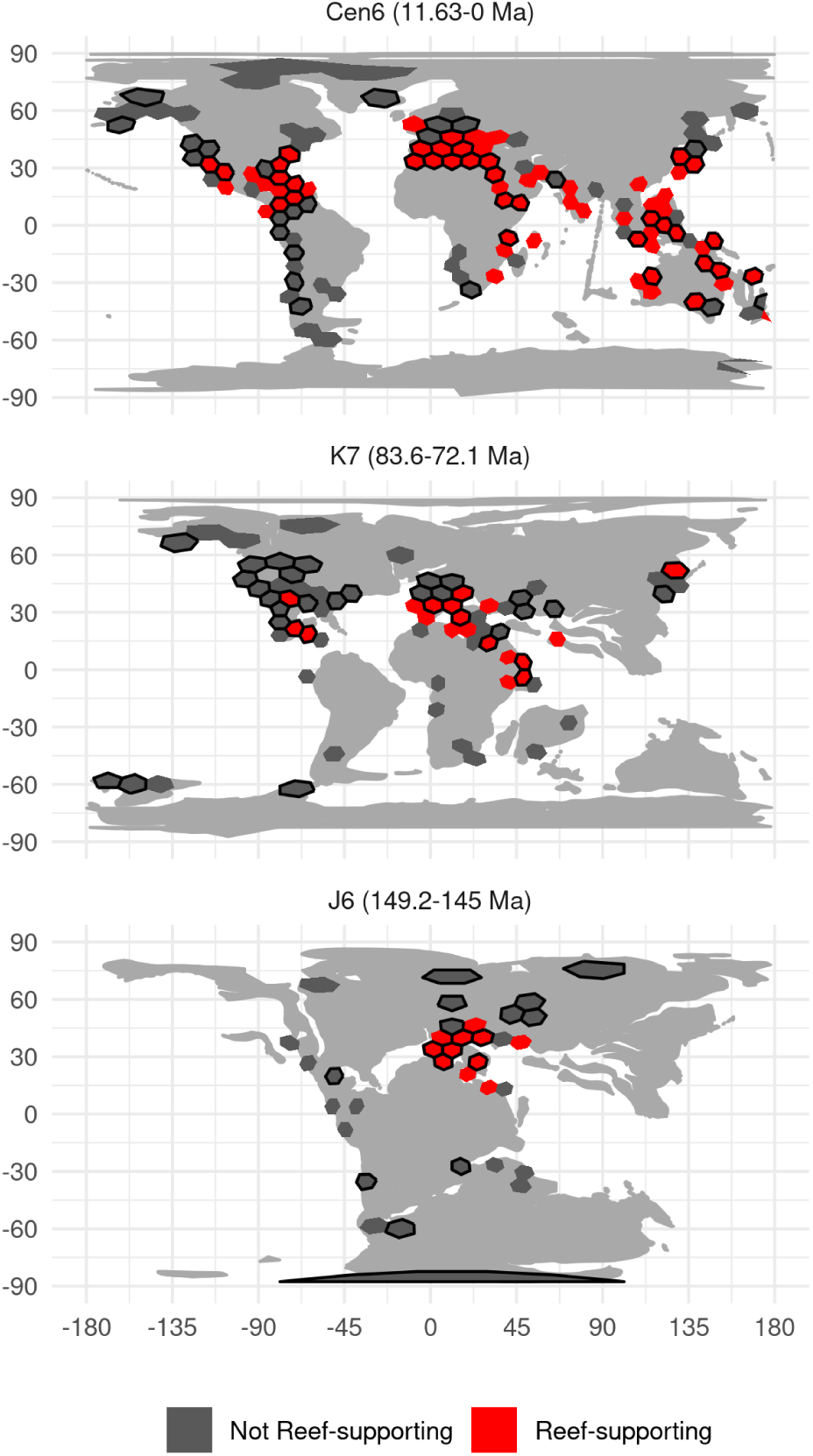
Paleogeographic distributions of reef-supporting (red) and non-reef-supporting (gray) regions (equal-area hexagonal/pentagonal grid cells with 1000 km spacings) for three of the 49 equal-length time intervals analyzed (Cen6, K7 and J6). Black borders denote grid cells that meet our quality criteria [i.e., cells containing more than 10 collections, 5 references, and a multiton ratio (*33*)] greater than 0.3), while those without black borders are those that do not. Cen6 (11.62 – 0.0117 Ma) = Messinian, Tortonian, Zanclean, Piacenzian, Gelasian, Calabrian, Middle Pleistocene, Late Pleistocene; K7 (83.6 – 72.1 Ma) = Campanian; J6 (152.1 – 145 Ma) = Tithonian. Paleomaps from the PALEOMAP project (*34*). See Fig. S1 for maps of all equal-length bins.

For counts of genera per fossil collections, we analyzed diversity patterns for reefal and non-reefal environments (facies). However, we classified grid cells and formations as either a “reef-supporting region” or a “non-reef-supporting region” based on the presence or absence of reefal collections [Fig. 1 and fig. S1; (*19*)]. Data for reefs is substantially less abundant than for most other facies (fig. S2), consistent with the observation that modern-day reefs account for only 0.1% of the Earth’s surface by area (*1*), and this imposes limits on our ability to infer regional-scale diversity patterns for reefal and non-reefal facies separately through time. Classifying grid cells based on the presence or absence of reefs is valid if diversity within reef-supporting regions is high compared to that of other regions that do not support reefs, independent (or partially independent) of the other facies represented by individual collections within those regions. We tested this hypothesis by quantifying diversity patterns separately for 1) reefal collections within reef-supporting grid cells (i.e., cells that have yielded collections representing reefal facies); 2) non-reefal collections within reef-supporting grid cells; and 3) non-reefal collections in non-reef-supporting cells (i.e., grid cells that have not yielded reefal collections). We held sampling intensity within grid cells constant by subsampling to equal counts of collections, using quotas of 20 and 40. We found that non-reefal collections from reef-supporting cells have similar mean diversities to those of reefal collections, and higher diversities than those of cells that lack reefs (fig. S3, table S1), supporting the designation of a cell as “reef-supporting” if containing any reefal facies. Various hypotheses may explain why non-reefal environments hosted higher diversity in reef-supporting regions than elsewhere, including environmental heterogeneity (*7*), conducive climate [for example, tropical coastal zones are more diverse than temperate zones today, even outside of reefs (*1*)], or because reefs are net exporters of species to surrounding environments (*7*).

### Patterns of diversity among reef-supporting and non-reef-supporting regions

We find that reef-supporting regions hosted between two- to three-fold higher levels of diversity on average than non-reef-supporting regions throughout the Phanerozoic, depending on grid cell size, pre- or post-Cretaceous–Paleogene time interval, and sifting criteria (figs. 2A and S4; for Wilcox tests of differences and effect sizes quantified as *r* values, see tables S2 and S3; additional spatial scales are shown in fig. S5). This is despite reef-supporting regions being less numerous and having a more restricted spatial distribution compared to non-reefal regions: non-reef-supporting cells outnumbered reef-supporting cells through most of the Phanerozoic (Fig. 2C), although counts of reef-supporting cells increase towards the present and marginally exceed non-reef-supporting cells in the most recent time bin.

**Fig. 2:**
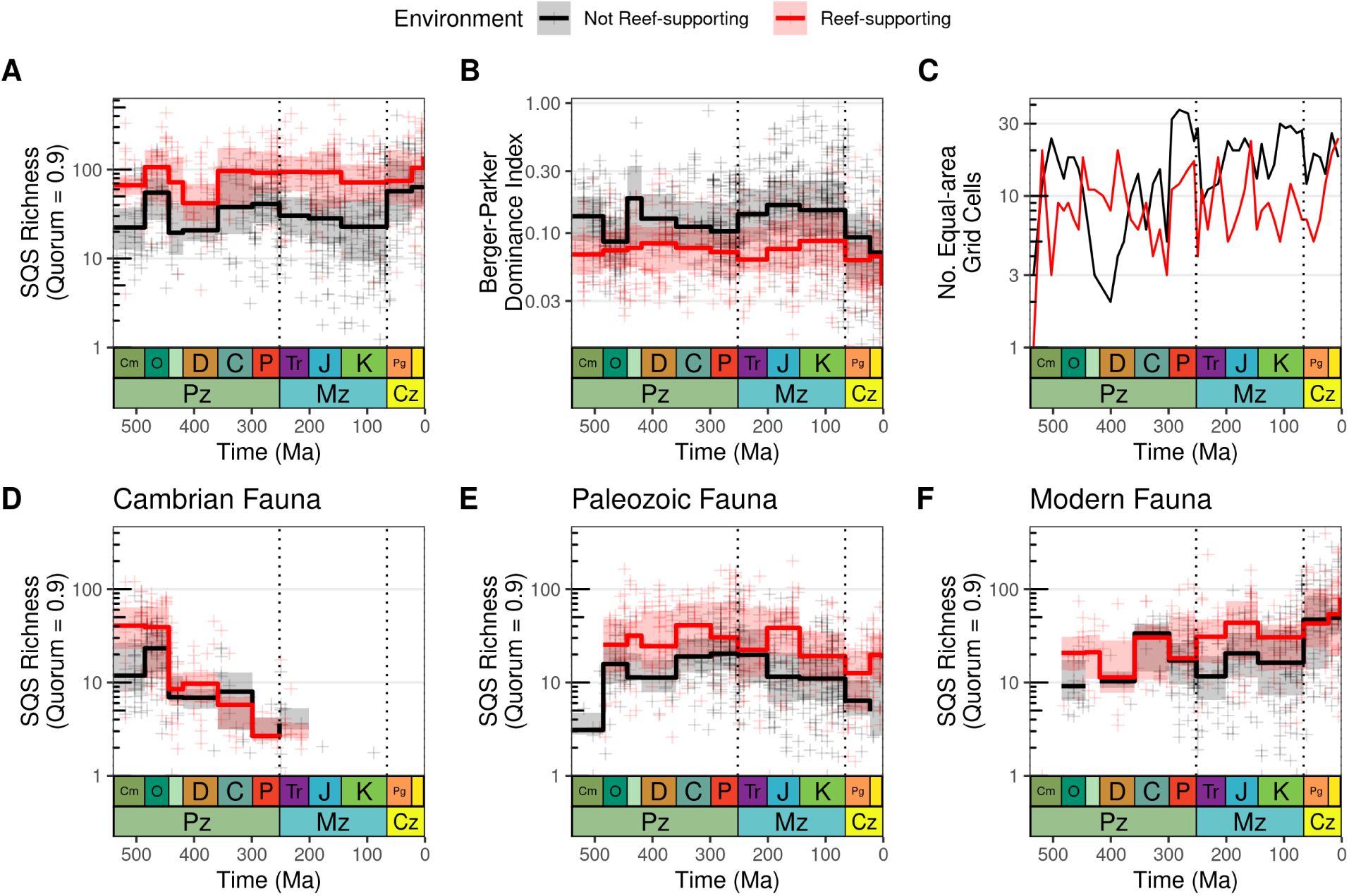
Differences in Phanerozoic marine invertebrate animal diversity patterns between reef-supporting (red) and non-reef-supporting (black) regions (equal-area hexagonal/pentagonal grid cells with spacings of 1000 km for all panels in this figure). Excluding collections explicitly identified as representing unlithified or poorly-lithified- and-sieved deposits, and also excluding collections that have no information about lithification style (see Fig. S7 for patterns using other sifting criteria). Dotted lines represent boundaries between geological eras. Note logarithmic y-axes. For panels A–B and D–F, crosses represent SQS diversity estimates for individual grid cell regions, while trend lines represent medians and interquartile ranges of regional diversity for geological periods. (A) Spatially-standardized Phanerozoic marine animal diversity, contrasting patterns for reef-supporting and non-reef-supporting regions. Note that in reef-supporting regions, levels of diversity have been broadly similar since the Ordovician, with no evidence for long-term, secular trends. In non-reef-supporting regions, by contrast, levels of diversity were similar from the Ordovician to the latest Cretaceous, when diversity rose fairly rapidly to a new, higher level that was sustained through the Cenozoic. However, this K/Pg increase is strongly associated with gastropods and unlithified sediments (see fig. S6). (B) Evenness, estimated using Berger-Parker dominance index (*35*), in reef-supporting and non-reef-supporting grid cells. (C) Counts of reef-supporting and non-reef-supporting cells through the Phanerozoic, using equal-length time bins. Panels (D–F) show patterns for Sepkoski’s evolutionary faunas. (D) Cambrian Fauna (Trilobita, Linguliformea, Graptolithina, Conodonta); (E) Modern Fauna (Anthozoa, Ostracoda, Rhynchonelliformea, Cephalopoda, Crinoidea); (F) Modern Fauna (Bryozoa, Bivalvia, Gastropoda, Echinoidea, Chondrichthyes).

Diversity within reef-supporting regions most likely reached modern levels by the Ordovician and did not experience substantial step-changes or sustained increases over the remainder of the Phanerozoic (Fig. 2A). Average levels of diversity in non-reef-supporting regions increased across the K/Pg boundary to approximately twice those attained throughout most of the preceding Phanerozoic, although a peak of comparable magnitude was reached earlier in the Ordovician (figs. S6A and 2A; see tables S3 and S4 for effect sizes). As a result, diversity within reef-supporting regions was approximately three times higher than non-reef-supporting regions on average during the Paleozoic–Mesozoic, but only approximately twice as high during the Cenozoic (table S4. It is possible that this increase may be exaggerated by the presence of less-well-lithified deposits in the dataset: only a modest increase is seen when analyzing lithified data only, but a pronounced increase is seen when including all data regardless of lithification style (fig. S7; table S4). The Cenozoic radiation of gastropods may also play a role in the observed increase (see below). Wilcoxon statistical tests show a highly-significant increase in diversity for non-reef-supporting regions from the Paleozoic/Mesozoic into the Cenozoic when including collections that lack metadata on lithification, but this is weaker when lithified-only data are analyzed (fig. S6A; table S4).

Reef-supporting regions are significantly more diverse than non-reef-supporting regions even at low paleolatitudes, demonstrating that this effect is not due solely to higher diversity in low-latitude tropical regions (Fig. 3, and figs. S8 and S9). The finding that reef-supporting regions were more diverse than non-reef-supporting regions is also consistent regardless of grid cell size choice, but differences become more pronounced as cell sizes increase (fig. S5; tables S2 and S3), suggesting that reef-supporting regions attain high diversity in part through elevated beta diversity. Nevertheless, even counts of genera per collection and per formation exhibit similar trends to those observed in reef-supporting vs non-reef-supporting cells, meaning that reefal collections and reef-supporting formations are more diverse than non-reefal collections and non-reef-supporting formations, and both are non-trending when only lithified deposits are analyzed (figs. S10 and S11).

**Fig. 3:**
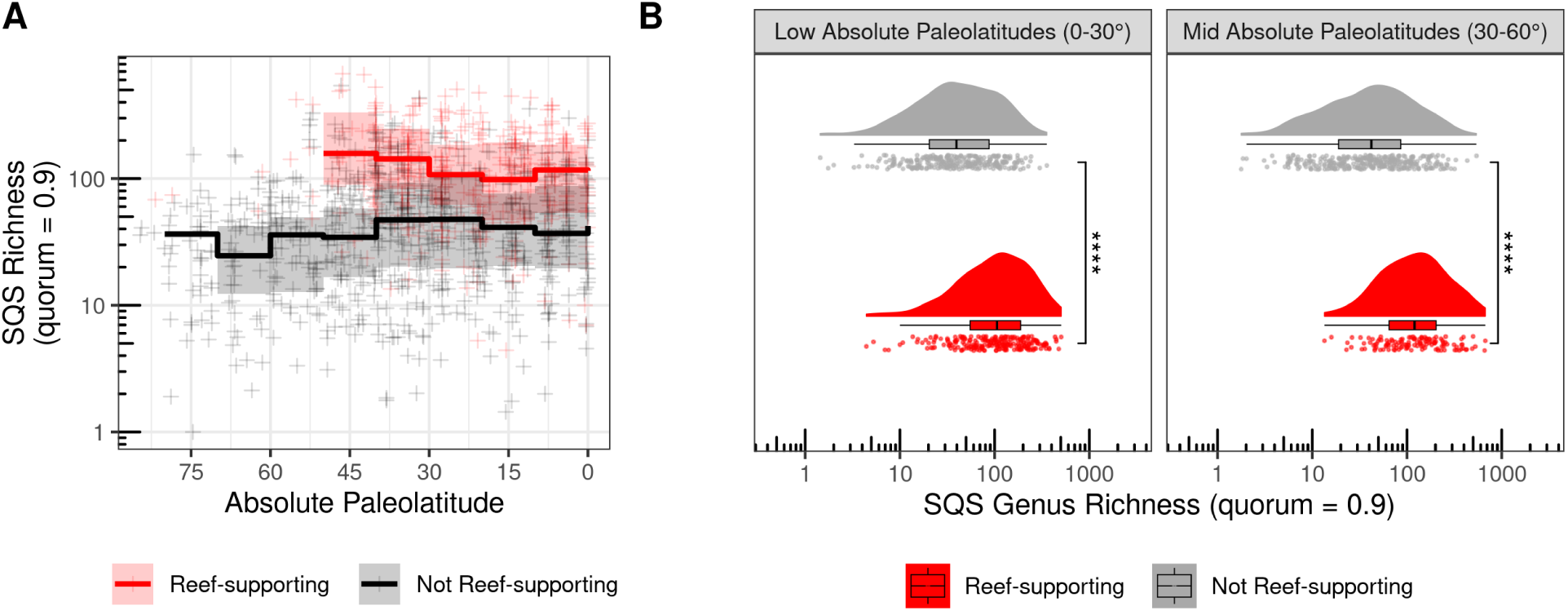
The effect of paleolatitude on diversity within reef-supporting and non-reef-supporting regions. (A) Latitudinal gradients of Phanerozoic marine invertebrate animal diversity, showing separate trends across paleolatitudes for reef-supporting (red) and non-reef-supporting (black) regions (equal-area hexagonal/pentagonal grid cells with spacings of 100 km, 500 km, 1000 km and 2000 km). Crosses represent SQS diversity estimates for individual grid cell regions, while trend lines represent medians and interquartile ranges of regional diversity for 10 degree paleolatitudinal bins. PaleoDB fossil collections that had been explicitly identified as representing unlithified or poorly-lithified-and-sieved deposits were excluded. Note logarithmic y-axis. Diversity is significantly higher in reef-supporting regions than in non-reef-supporting regions, even at low paleolatitudes, indicating that this effect is not solely driven by higher diversity in all tropical low-paleolatitude regions. (B) Distributions of SQS diversity (quorum = 0.9) within reef-supporting (red) and non-reef-supporting (gray) regions (equal-area hexagonal/pentagonal grid cells with 100 km, 500 km, 1000 km and 2000 km spacings) between low (0–30°) and mid (30–60°) absolute paleolatitude zones. Diversity is consistently significantly higher in reef-supporting regions than in non-reef-supporting regions, regardless of paleolatitude zones. Each plot shows three ways of visualizing the data: individual data points with jitter, a boxplot (line denotes the median value, the hinges of the box correspond to the interquartile range, and whiskers extend from the hinges to the smallest/largest values at most 1.5 * IQR of the hinge), and a density plot using a Gaussian kernel with a smoothing bandwidth of 1. Statistical significance for Wilcoxon tests between groups is indicated by either no text (p ¿ 0.05), * (p ≤ 0.05), ** (p ≤ 0.01), *** (p ≤ 0.001) or **** (p ≤ 0.0001).

Higher overall diversity in reef-supporting regions does not appear to result from dramatically higher richness of individual clades (fig. S12), but rather from the summation of modest differences in diversity across a wide range of major groups, such as brachiopods, bivalves, and gastropods, with a greater number of higher taxa in each reef-supporting region (fig. S13). Although reef-supporting regions host markedly higher diversity of clades such as corals, sponges and sea-lilies, the overall diversity dynamics of other groups are superficially similar, and even non-reef-building organisms have higher diversity within reef-supporting regions (fig. S12). Parsing out diversity patterns for reef-supporting regions according to the reef-building organisms that are present within each cell (Fig. 4 and fig. S14) shows that reef-supporting cells containing corals are the most consistently diverse (Ordovician–present), followed by those containing microbial reef-builders (Cambrian–Cretaceous), while stromatoporoids were important in the Paleozoic and Jurassic–Cretaceous. Reef-supporting regions also have systematically more even occurrence-frequency distributions than non-reef-supporting regions as measured by the Berger-Parker dominance index (Fig. 2B), possibly reflecting the different structures of these ecological communities. Non-reef-supporting regions show a substantial increase in evenness over the latest-Cretaceous/Cenozoic interval, consistent with ref (*21*) (Fig. 2B).

**Fig. 4:**
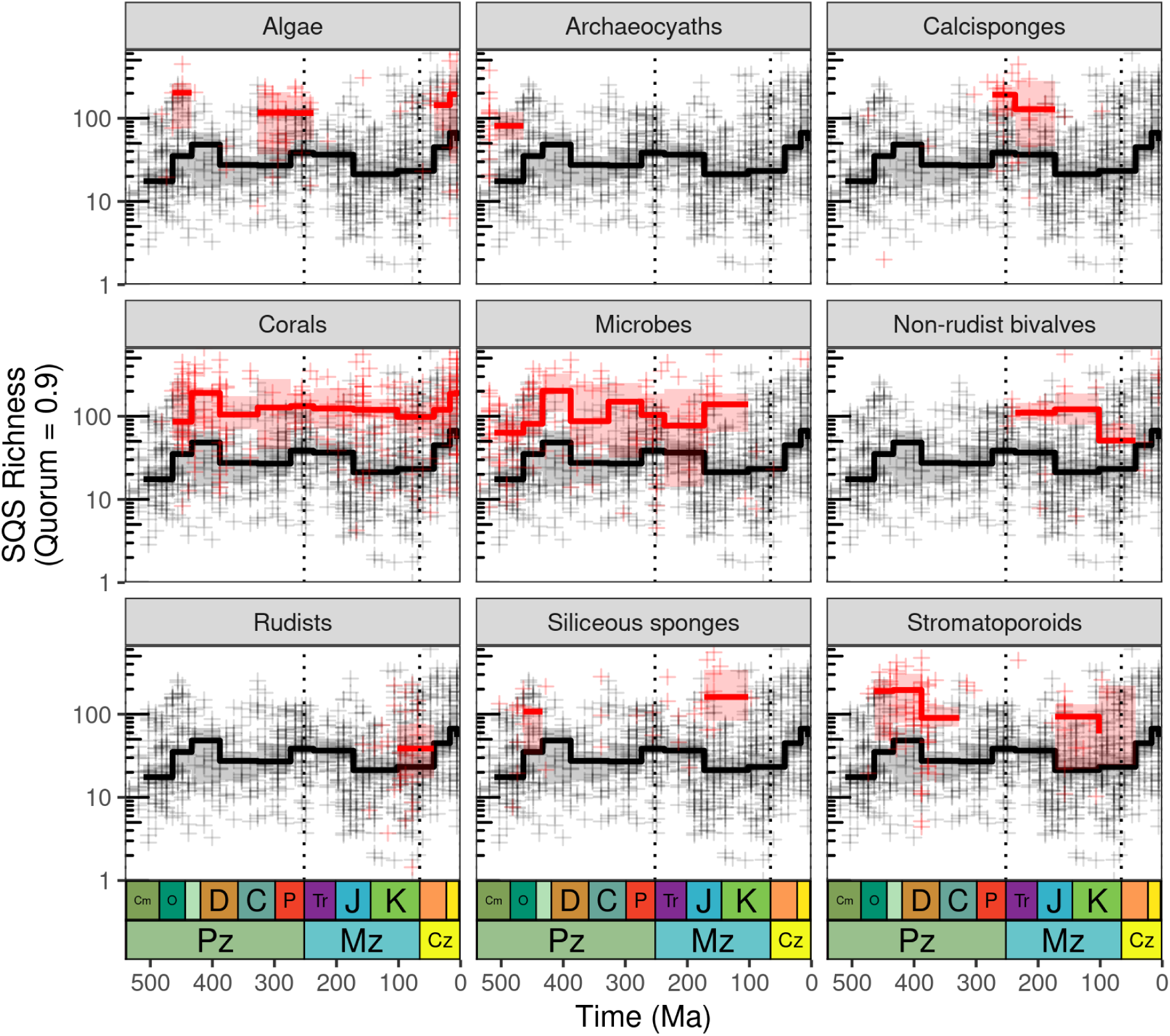
Differences in diversity patterns (SQS, quorum = 0.9) for marine invertebrate animals between reef-supporting (red) and non-reef-supporting (black) regions (equal-area hexagonal/pentagonal grid cells with 1000 km spacings), showing patterns for major kinds of reef-building organisms (see fig. S14 for additional reef-building organisms). Reef-supporting cells were identified as being associated with each particular kind of reef-building organism using the palaeocoordinates of reef sites listed in the PARED PaleoReefs database (*36*) (grid cells can appear in more than one panel if they contain more than one type of reef-building organism). Note logarithmic y-axes. Crosses represent SQS diversity estimates for individual grid cell regions, while trend lines represent medians and interquartile ranges of regional diversity for geological periods. Dashed lines represent boundaries between geological eras. PaleoDB collection data excludes those identified as unlithified and poorly-lithified-and-sieved deposits, but includes collections with no information on lithification style.

Across all three of Sepkoski’s ‘evolutionary faunas’ (*22*), diversity is higher in reef-supporting than in non-reef-supporting regions (Fig. 2D–F). However, the diversity dynamics of Sepkoski’s ‘Modern Fauna’ (*22, 23*), which includes groups such as gastropods, bivalves, and echinoids, differ substantially between reef-supporting and non-reef-supporting regions (Fig. 2F; see also fig. S12). The Modern Fauna became an important component of reef-supporting regions during the Paleozoic, showing only a weakly-increasing mean diversity trajectory from the Carboniferous (358.9 Ma) onward (Fig. 2F). In contrast, the Modern Fauna was less diverse in non-reef-supporting regions during the Mesozoic (with the exception of the Carboniferous), but its diversity in these regions increased steadily until the late Mesozoic, after which it increased abruptly around the K/Pg. The Paleozoic Fauna, primarily comprising articulate brachiopods, crinoids, corals, cephalopods, and bryozoans, had higher average diversity in reef-supporting regions than in non-reef-supporting regions (Fig. 2E). Overall diversity levels for the Paleozoic Fauna in both reef- and non-reef-supporting regions, however, remained more or less stable through the Paleozoic, followed by a gradual decrease through the Mesozoic and Cenozoic and a marked drop after the end-Cretaceous mass extinction, especially in non-reef-supporting regions. Despite these differential faunal dynamics, however, diversity for marine animals as a whole fluctuated around two different, but relatively unchanging, average levels in both reef-supporting and non-reef-supporting regions over much of the Phanerozoic (Fig. 2A). Across all three of Sepkoski’s ‘evolutionary faunas’ (*22*), diversity is higher in reef-supporting than in non-reef-supporting regions (Fig. 2D–F). However, the diversity dynamics of Sepkoski’s ‘Modern Fauna’ (*22, 23*), which includes groups such as gastropods, bivalves, and echinoids, differ substantially between reef-supporting and non-reef-supporting regions (Fig. 2F; see also fig. S12). The Modern Fauna became an important component of reef-supporting regions during the Paleozoic, showing only a weakly-increasing mean diversity trajectory from the Carboniferous (358.9 Ma) onward (Fig. 2F). The increase in diversity of the Modern Fauna over the K/Pg boundary interval is markedly more pronounced in non-reefal regions (Fig. 2F). The Paleozoic Fauna, primarily comprising articulate brachiopods, crinoids, corals, cephalopods, and bryozoans, had higher average diversity in reef-supporting regions than in non-reef-supporting regions (Fig. 2E). Overall diversity levels for the Paleozoic Fauna in both reef- and non-reef-supporting regions, however, remained more or less stable through the Paleozoic, followed by a gradual decrease through the Mesozoic and Cenozoic and a marked drop after the end-Cretaceous mass extinction, especially in non-reef-supporting regions. Despite these differential faunal dynamics, however, diversity for marine animals as a whole fluctuated around two different, but relatively unchanging, average levels in both reef-supporting and non-reef-supporting regions over much of the Phanerozoic (Fig. 2A).

The increase in diversity of the Modern Fauna over the K/Pg boundary interval is markedly more pronounced in non-reefal regions (Fig. 2F). Much of this shift is driven by increases in gastropod diversity, which were proportionally greater in non-reefal than in reefal regions from the Maastrichtian onward (figs. S6B and S12; table S5). The effect of this radiation on diversity patterns, however, is difficult to disentangle from the effects of changing lithification styles (fig. S15), because the availability of lithified deposits diminished over this interval (figs. S16 and S6A). A large fraction of extant gastropod richness is concentrated in the very small “micromollusc” size range, which would be more easily recovered from unlithified sediments (*24, 25*).

We show that levels of richness were non-directional in both reef-supporting and non-reef-supporting regions over an interval of 400 million years. This contradicts classical hypotheses of directional long-term increasing diversity throughout geological time [e.g., refs (*22, 26*)]. Our finding that reef-supporting regions supported high levels of diversity that did not change systematically over long intervals of geological time complements existing characterizations of boom-and-bust patterns of reef crises (*27*) over shorter time intervals, prompted by poorly-understood threshold effects. Analyses on a shorter temporal scale reveal the fluctuations in reef-supporting diversity in step with these crises (figs. S17 and S18), but when observed over tens to hundreds of millions of years, these environments fluctuated around a relatively unchanging average level of diversity (Fig. 2A).

## Conclusions

Our study demonstrates that reef-supporting regions have hosted a disproportionate fraction of marine biodiversity throughout the Phanerozoic, despite profound changes in the taxonomic composition of reef-builders over this interval (*28*). This result is consistent with previous studies that found statistical evidence for the presence or prevalence of reefs being associated with high levels of diversity, either at notionally “global” (*8*) or regional scales (*10*). These findings may also be consistent with previous work showing that reefs and shallow-water tropical environments during the Phanerozoic were sites of high origination and acted as cradles of diversity that exported species to other environments (*7*). Foundational studies of paleobiology suggest that the groups comprising the Modern Fauna became highly diverse in wider marine ecosystems only after the Paleozoic (*22*). However, we show that they were an important component of reef-supporting regions much earlier, during the Paleozoic, suggesting an important role for reefs as cradles of higher taxonomic composition throughout the evolutionary history of animals. As hotspots of diversity and potential net exporters of species to other marine environments, reefal instability into the future (*29–32*) could therefore restructure marine ecosystems with major consequences for marine biodiversity over extended geological timescales.

## Acknowledgments

We thank Shanan Peters, Nadia Santodomingo, and Norman MacLeod for discussion. We thank the founders and contributors to the Paleobiology Database. This is Paleobiology Database publication number XXXX.

## Author contributions

Conceptualization: RAC, ES, RBJB Methodology: RAC, RBJB, ES

Investigation: RAC Visualization: RAC

Funding acquisition: ES, RBJB, RAC

Project administration: ES, RBJB, RAC

Writing – original draft: RAC

Writing – review & editing: RAC, ES, RBJB, WK

## Data and materials availability

(The data and analysis code will be available on Dryad/FigShare/Zenodo upon publication.)

## Supplementary Materials

### Materials and Methods

#### Occurrence data downloads

We reconstructed patterns of local- and regional-scale diversity for Phanerozoic marine invertebrates and major sub-groups using occurrence data downloaded from the Paleobiology Database (*18*). Genus-level occurrence data for the taxon set Animalia excluding Chordata from marine environments were downloaded from the Paleobiology Database on 11 July 2024, using the PaleoDB API URL “https://paleobiodb.org/data1.2/occs/list.csv?base_name=AnimaliâChordata&idtype=latest&pres=regular&envtype=marine&idreso=lump_gensub&idqual=genus_certain&taxon_status=valid&show=attr,classext,subgenus,abund,ecospace,taphonomy,ctaph,etbasis, pres,coll,coords,loc,paleoloc,time,timebins,stratext,lithext,env,geo,methods,resgroup,ref, refattr,ent,entname,crmod”. Subgenera were elevated to their respective genera. In order to visualize patterns within groups and to reconstruct patterns within Sepkoski’s three evolutionary faunas (*14*), we additionally defined taxon sets for major groups of marine animals, including Anthozoa, Bivalvia, Brachiopoda, Cephalopoda, Crinoidea, Echinoidea, Gastropoda, Porifera, and Trilobita, using downloads of PaleoDB occurrences numbers. Sepkoski’s evolutionary faunas comprise the Cambrian Fauna (Trilobita, Linguliformea, Graptolithina and Conodonta); the Paleozoic Fauna (Anthozoa, Ostracoda, Rhynchonelliformea, Cephalopoda and Crinoidea), and the Modern Fauna (Bryozoa, Bivalvia, Gastropoda, Echinoidea and Chondrichthyes). Prior to cleaning and binning, there were 800,928 occurrences.

#### Occurrence data cleaning

Occurrence data were cleaned following standard practices for removing unsuitable occurrences [e.g., refs (*8, 10*)]. We excluded (1) uncertain identifications (i.e., retaining only valid taxa), (2) collections with geographic scale listed as “basin” or stratigraphic scale listed as “group”, (3) fossils preserving soft parts, (4) coprolites and traces (i.e., retaining only body fossils with preservation mode given as “regular”), and (5) terrestrial and freshwater taxa.

To more fully control for biases arising from systematic changes in the mode of lithification of fossil-bearing sediments through the Phanerozoic, we restrict our focal analyses to collections that explicitly represent lithified deposits, or poorly-lithified deposits that have not been sieved (i.e., excluding unlithified and poorly-lithified and sieved deposits). Lithification biases arise because unlithified and poorly-lithified deposits are more prevalent in the fossil record from the Late Cretaceous through to the Recent [refs (*37, 38*); fig. S16]. Counts of taxa from fossil deposits that are unlithified or poorly-lithified are inflated relative to those from lithified sediments because they facilitate easier extraction of fossil specimens, especially those that are small (*37, 39–41*). Past studies of Phanerozoic marine animal diversity [e.g., refs (*8, 10, 15, 16, 42*)] have excluded fossil data identified as explicitly originating from deposits that are either unlithified, or poorly-lithified and sieved. However, many collections in the PaleoDB lack information about lithification style (n = 53,516, or 37.2% of cleaned data). Past studies typically have not excluded collections lacking lithification data, but our analyses (see below) suggest that many of these untagged collections likely derive from unlithified sediments. Because of this, it is not possible to fully control for lithification biases when including collections that lack information on lithification style.

Therefore, all of our focal analyses exclude unlithified or poorly-lithified and sieved data, as per previous studies, but additionally exclude collections lacking information on lithification style. However, we also present results for multiple additional sifting criteria, including only excluding data that had been specifically tagged as being unlithified or poorly-lithified and sieved, as per previous studied, and results are similar (fig. S7). Our full set of sifting criteria comprise: (1) all lithification styles, including collections lacking information on lithification; (2) excluding collections representing both unlithified and poorly-lithified-and-sieved deposits, but retaining collections lacking information on lithification style [the default sifting criteria used by previous studies of Phanerozoic marine animal diversity; e.g., refs (*10, 15, 16*)]; (3) excluding collections representing both unlithified and poorly-lithified-and-sieved deposits, and additionally excluding collections lacking information on lithification style; (4) collections explicitly identified as representing lithified deposits; (5) unlithified or poorly-lithified collections only; and (6) collections lacking information on lithification style only.

After removing unsuitable occurrences and binning into approximately equal-length time bins (see below; table S7), the data comprised 143,773 collections containing 699,642 occurrences, representing 30,209 genera (*18*).

#### Time binning of occurrence data

Occurrence data were binned into approximately equal-length time bins to control for the high variability in geological stage durations, especially towards the present-day [table S7; this practice follows past studies, such as refs (*8, 10*)]. The cleaned and binned occurrence data comprised 143,773 collections containing 699,642 occurrences, representing 30,209 genera (*18, 19*).

#### Spatial standardization

Paleocoordinates for fossil collections were rotated to the map age closest to the midpoint of each time bin using the R package chronosphere [version 0.6.1, online reconstruction method; ref (*43*)] with the PALEOMAP paleogeographic model of C. Scotese (*34*). We standardized the extent of spatial sampling by binning fossil localities into equal-area hexagonal/pentagonal grid cells (*19*) with 100 km, 500 km, 1000 km, and 2000 km spacings (Fig. 1) using the dggridR R package version 2.0.3 (*44*). These grid-cell spacings were chosen to balance regional extent versus the need for sufficient sampling, and are comparable to regional spatial samples used in our previous work (*10*).

#### Calculations of regional diversity and other variables

For each grid cell, we calculated sampling-standardized genus richness using Shareholder Quorum Subsampling [SQS (*8, 15*), also known as coverage-based rarefaction or CBR (*20, 45*)], at a quorum level (target sample coverage) of 0.9, using the R package iNEXT version 2.0.20 (*46*), which allows both drawing down samples (subsampling or interpolating) and extrapolating to equal coverage. Following (*20*), we discarded extrapolated estimates based on extrapolated samples that were more than twice the reference sample size. We excluded cells that did not meet certain data quality criteria indicating minimal sampling (our ‘quality criteria’). Cells were excluded if they contained fewer than 10 collections or had a multiton ratio [the proportion of species in a sample represented by at least two individuals (*33*)] of less than 0.3. This process eliminated grid cells that had spuriously high levels of sample completeness [measured by Good’s *u* (*47*)] resulting from estimation error due to small sample sizes (i.e., poorly-sampled grid cells that spuriously appeared to have high sample coverage by virtue of there being no or few singletons, despite very limited sampling).

We also calculated a range of other variables for each grid cell, including the Berger-Parker dominance index (*35*) as a measure of evenness, face-value (raw, or unstandardized) counts of genera, counts of fossil collections (the terminology used in the Paleobiology Database for samples of taxa from fossil localities that are well-circumscribed in time and space), counts of fossil occurrences (i.e., the confirmed presence of a taxon in a fossil collection), counts of references (i.e., publications associated with fossil occurrences), counts of geological formations, and counts of distinct paleocoordinate locations.

For Fig. 4, which parses reef-supporting and non-reef-supporting patterns according to the kinds of reef-building organisms present in each cell, reef-supporting cells were identified as being associated with particular kinds of reef-building organisms (using the palaeocoordinates of reef sites listed in the PARED PaleoReefs database (*36*) (note that grid cells can appear in more than one panel if they contain more than one type of reef-building organism).

#### Local and formation scale richness

To analyze patterns of genus richness at the local scale, we tallied face-value counts of genera per collection and per formation. Face-value counts at these spatial scales are heavily right-skewed [i.e., most collections contain only one or a small handful of taxa, but some collections are extraordinarily diverse, and represent a fairly complete snapshot of local-scale diversity (*4, 48*)]. To exclude uninformative collections and formations, we imposed quality criteria indicative of sampling level, based on counts of references, collections, and higher taxa (defined here as Mollusca, Arthropoda, Echinodermata, Cephalopoda, Anthozoa, Chordata *sans* Tetrapoda, Bryozoa, Porifera, Graptolithina, and Annelida): we retained collections with at least five references per collection or four higher taxa, and formations with at least 20 collections, five references or four higher taxa.

#### Classification of reef-supporting and non-reef-supporting grid cells

We classified cells as reef-supporting if they contained at least one fossil collection that represented reefal facies in the Paleobiology Database occurrence download. Reefal environmental facies were identified using the ‘environment’ field and the values “reef, buildup or bioherm”, “slope/ramp reef”, “perireef or subreef”, “basin reef”, “platform/shelf-margin reef”, and “intrashelf/intraplatform reef”, while non-reefal facies comprised all other terms (“offshore”, “peritidal”, “carbonate indet.”, “offshore shelf”, “shallow subtidal indet.”, “lagoonal”, “sand shoal”, “transition zone/lower shoreface”, “deep subtidal shelf”, “offshore ramp”, “submarine fan”, “prodelta”, “shoreface”, “offshore indet.”, “deep subtidal indet.”, “open shallow subtidal”, “deltaic indet.”, “deep subtidal ramp”, “foreshore”, “interdistributary bay”, “delta plain”, “estuary/bay”, “coastal indet.”, “delta front”, “lagoonal/restricted shallow subtidal”, “paralic indet.”, “deep-water indet.”). If only imprecise environmental types were present in a cell (e.g., “marine indet.”), then that grid cell was not assigned to either reef-supporting or non-reef-supporting and was discarded.

We consider that the presence of a single reefal collection is sufficient to show that the geographic region encompassed by the cell was conducive to supporting reefal ecosystems, and would not lead to any ‘false positives’ in assigning cells to the reef-supporting category. Indeed, ‘false negatives’ are likely to be a more important source of error, due to cells that did in fact support reefal ecosystems in deep time, but which have not yielded reefal facies due to their restricted spatial distribution and thus lower likelihood of preservation. There is a linear increase in the log of SQS diversity with the number of reefal collections present in a cell (fig. S19B). There is also a strong linear relationship between the log of SQS diversity and the total number of collections present in a cell for both reef-supporting and non-reef-supporting regions, albeit with reef-supporting regions showing a steeper relationship than non-reef-supporting regions (fig. S19A). The proportion of reefal cells is a poor choice for assigning cells to reef-supporting or non-reef-supporting categories, not least because the number of total collections present in a cell declines as the proportion of reefal collections increases (fig. S19D), which causes the diversity of cells to decrease as the proportion of reefal collections increases (fig. S19C).

#### Statistical tests

To test for statistically-significant differences in diversity among groups (e.g., between reef-supporting and non-reef-supporting regions during the Pre- and Post-K/Pg time intervals, and within reef-supporting and non-reef-supporting regions across the Pre- and Post-K/Pg intervals), we conducted non-parametric Wilcoxon tests using the function wilcox test() in the R package ‘rstatix’ (version 0.7.0). To quantify effect sizes for differences between groups in Wilcoxon tests, we also computed the rank-biserial correlation (‘*r* value’) using the function wilcox effsize() in the R package ‘rstatix’, which is more appropriate than Cohen’s *d* (*49*) for this non-parametric test.

### Supplementary Text

End-Cretaceous to Cenozoic collections from non-reef-supporting regions that lack metadata on lithification style may include many collections with unlithified or poorly-lithified-and-sieved sediments (fig. S7). To control for this lithification bias (higher diversity in non-lithified deposits due to greater ease of extraction, especially for small specimens), it is necessary to exclude deposits that are unlithified or poorly-lithified and sieved. However, many PaleoDB collections lack information about lithification style, and it is likely that many of these (at least, those that date to the Late Cretaceous–Recent) represent unlithified or poorly-lithified deposits. When the ‘untagged’ data is included in the analysis, the increase in diversity in the Cenozoic is much more pronounced, especially in non-reef-supporting regions (fig. S6 and table S3). However, when the analysis is restricted to data that has been explicitly identified as originating from lithified deposits, the increase in the Cenozoic is substantially reduced in magnitude.

---

**Fig. S1:**
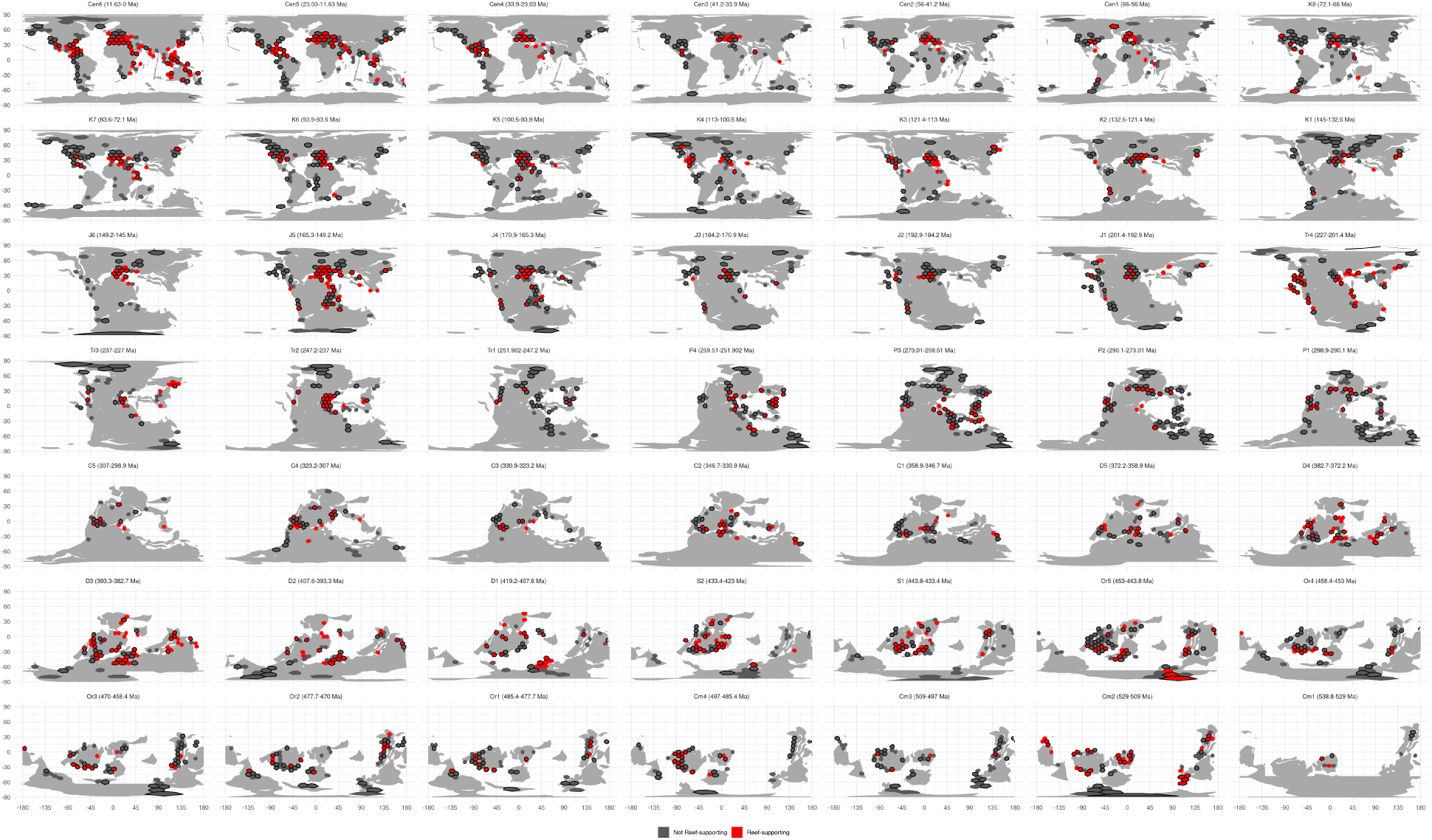
Paleogeographic distributions of reef-supporting (red) and non-reef-supporting (gray) regions (equal-area hexagonal/pentagonal grid cells with 1000 km spacings) for all of the equal-length time intervals analyzed (table S7). Black borders denote grid cells that meet our quality criteria, while those without black borders are those that do not. Paleomaps from the PALEOMAP project (*34*).

**Fig. S2:**
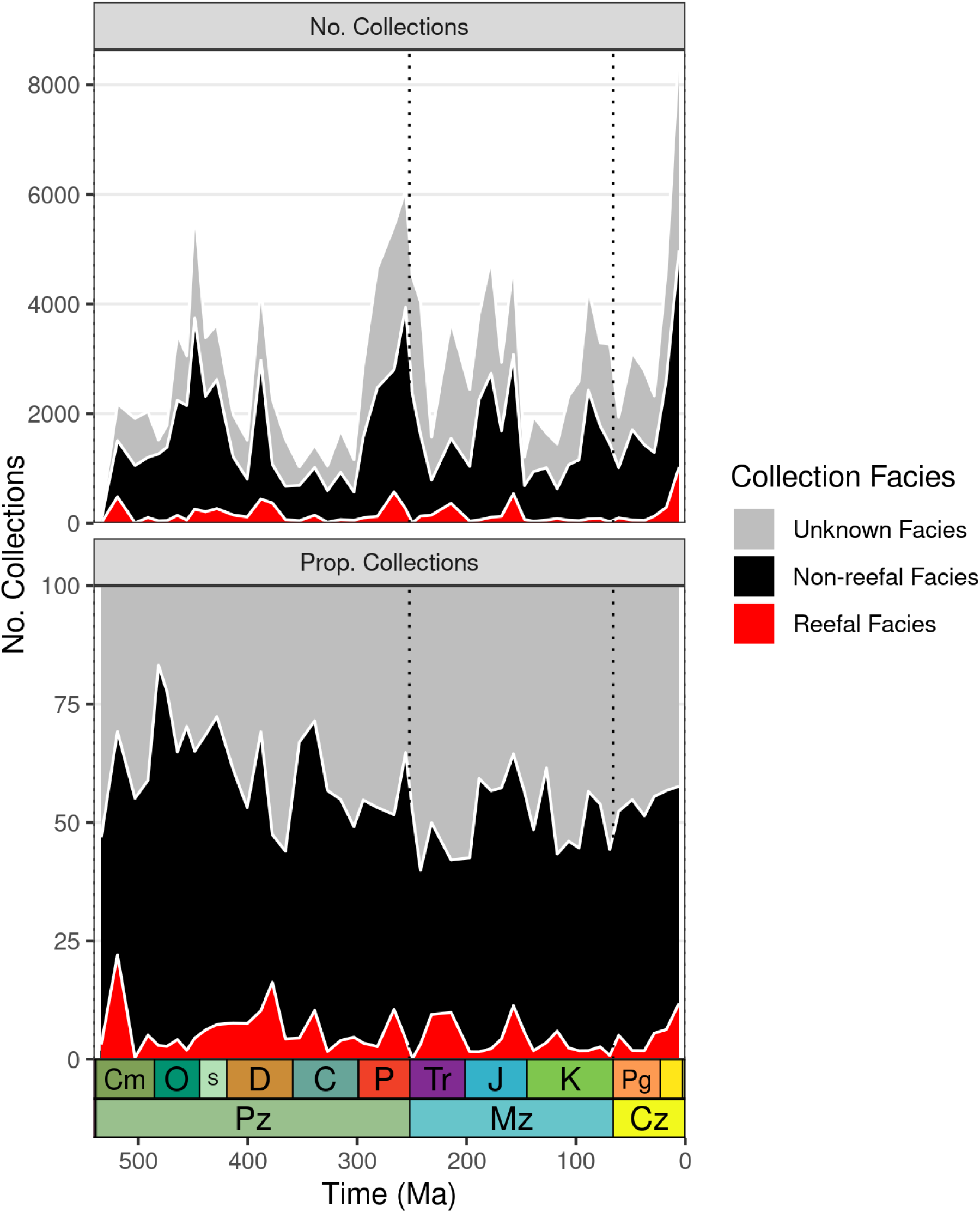
Counts of collections representing either reefal facies, non-reefal facies, or unknown facies (i.e., PaleoDB collections lacking information on environmental facies) through the Phanerozoic, using equal-length time bins.

**Fig. S3:**
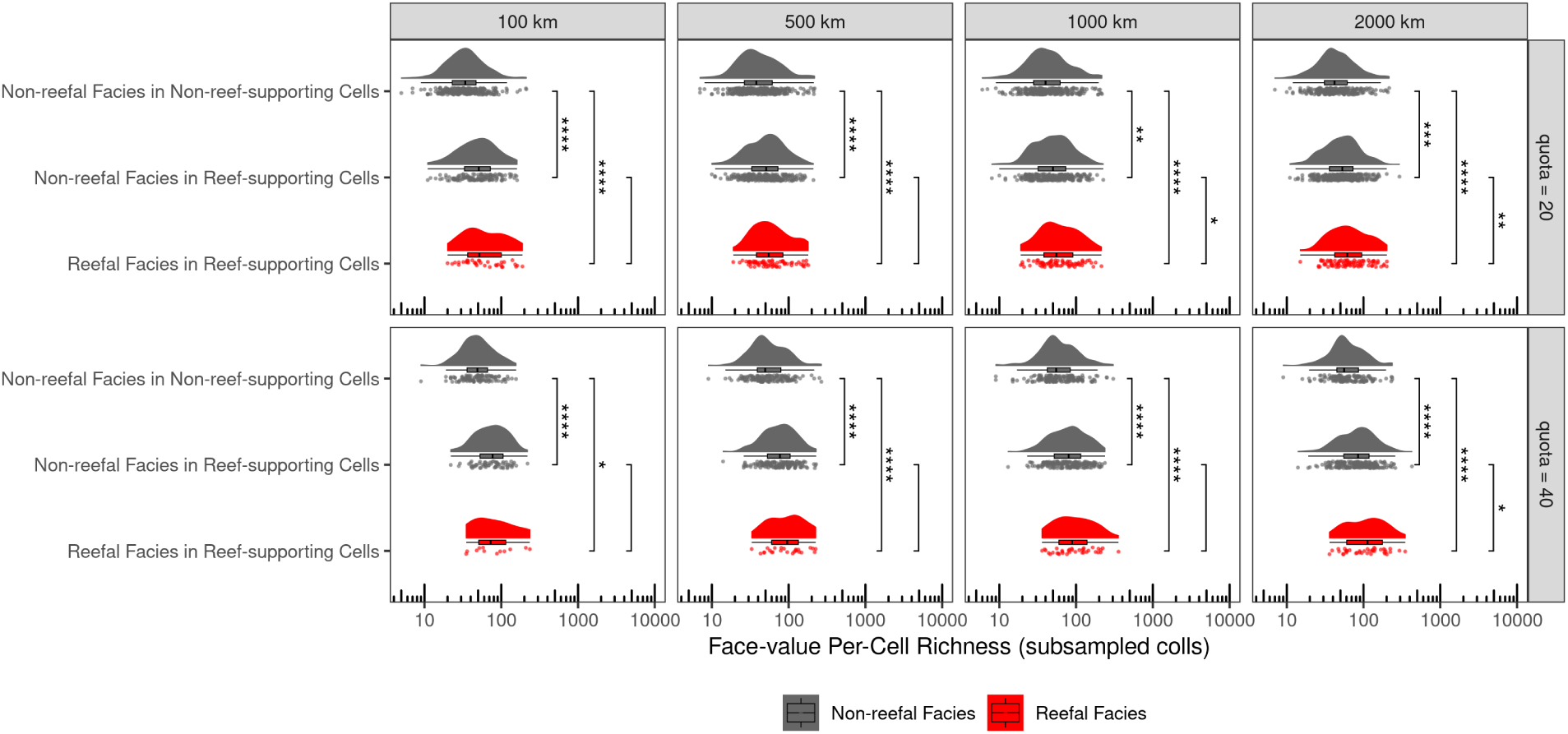
Distributions of diversity for Phanerozoic reef-supporting and non-reef-supporting regions (defined using equal-area grid cells of 100 km, 500 km and 1000 km) after subsampling to equal counts of reefal (red) or non-reefal (gray) collections at quotas of 20 and 40. These results show that when controlling for variation in sampling intensity within cells, reef-supporting cells host higher levels of diversity than non-reef-supporting cells, regardless of whether reefal facies or non-reefal facies are analyzed. Each plot shows three ways of visualizing the data: individual data points with jitter, a boxplot (line denotes the median value, the hinges of the box correspond to the interquartile range, and whiskers extend from the hinges to the smallest/largest values at most 1.5 * IQR of the hinge), and a density plot using a Gaussian kernel with a smoothing bandwidth of 1. Statistical significance for Wilcoxon tests between groups is indicated by either no text (p ¿ 0.05), * (p ≤ 0.05), ** (p ≤ 0.01), *** (p ≤ 0.001) or **** (p ≤ 0.0001).

**Fig. S4:**
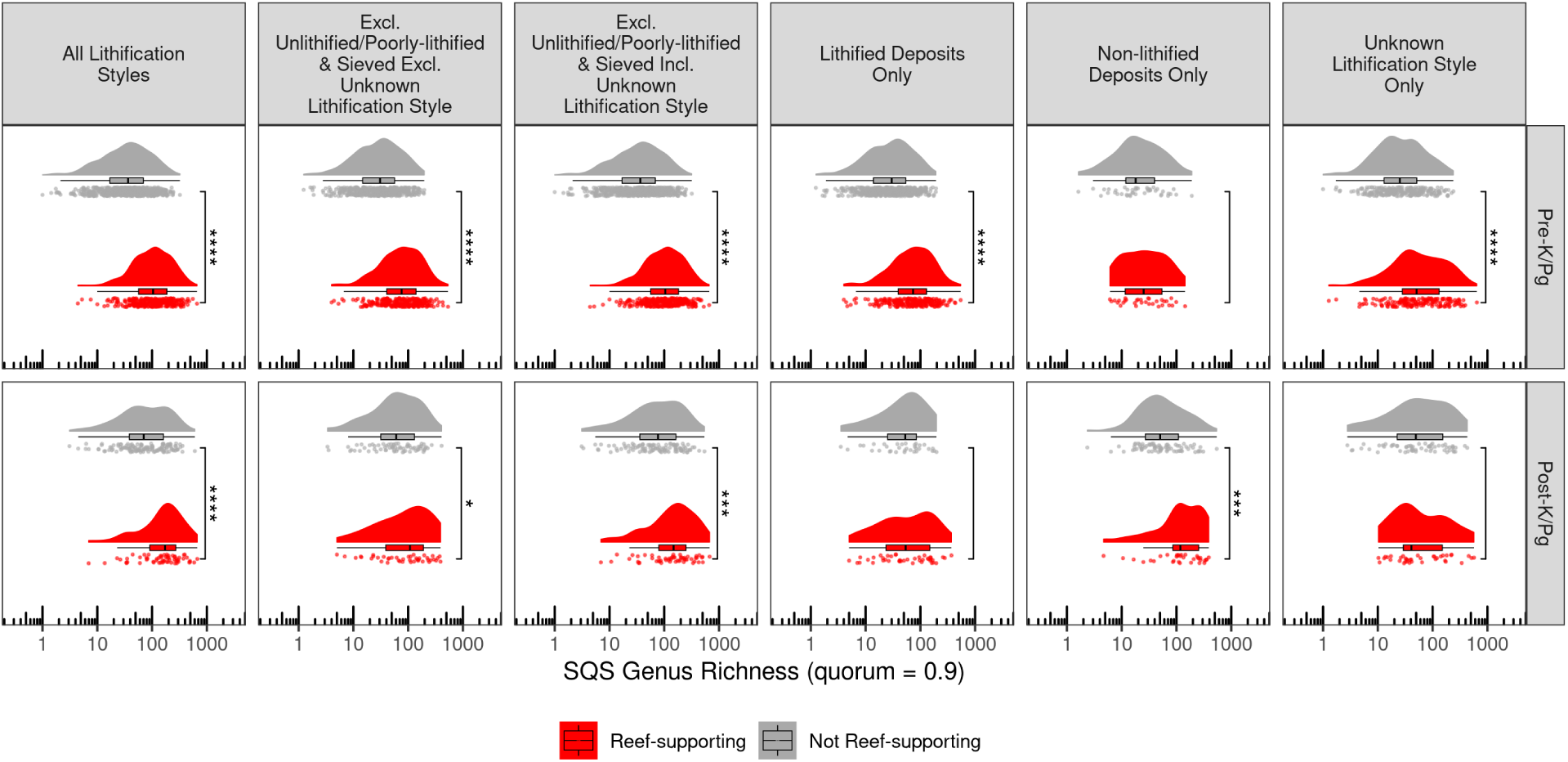
Distribution of diversity within reef-supporting (red) and non-reef-supporting (gray) regions (equal-area hexagonal/pentagonal grid cells with 1000 km spacings) in the Paleozoic–Mesozoic and Cenozoic. Each plot shows three ways of visualizing the data: individual data points with jitter, a boxplot (line denotes the median value, the hinges of the box correspond to the interquartile range, and whiskers extend from the hinges to the smallest/largest values at most 1.5 * IQR of the hinge), and a density plot using a Gaussian kernel with a smoothing bandwidth of 1. Statistical significance for Wilcoxon tests between groups is indicated by either no text (p ¿ 0.05), * (p ≤ 0.05), ** (p ≤ 0.01), *** (p ≤ 0.001) or **** (p ≤ 0.0001).

**Fig. S5:**
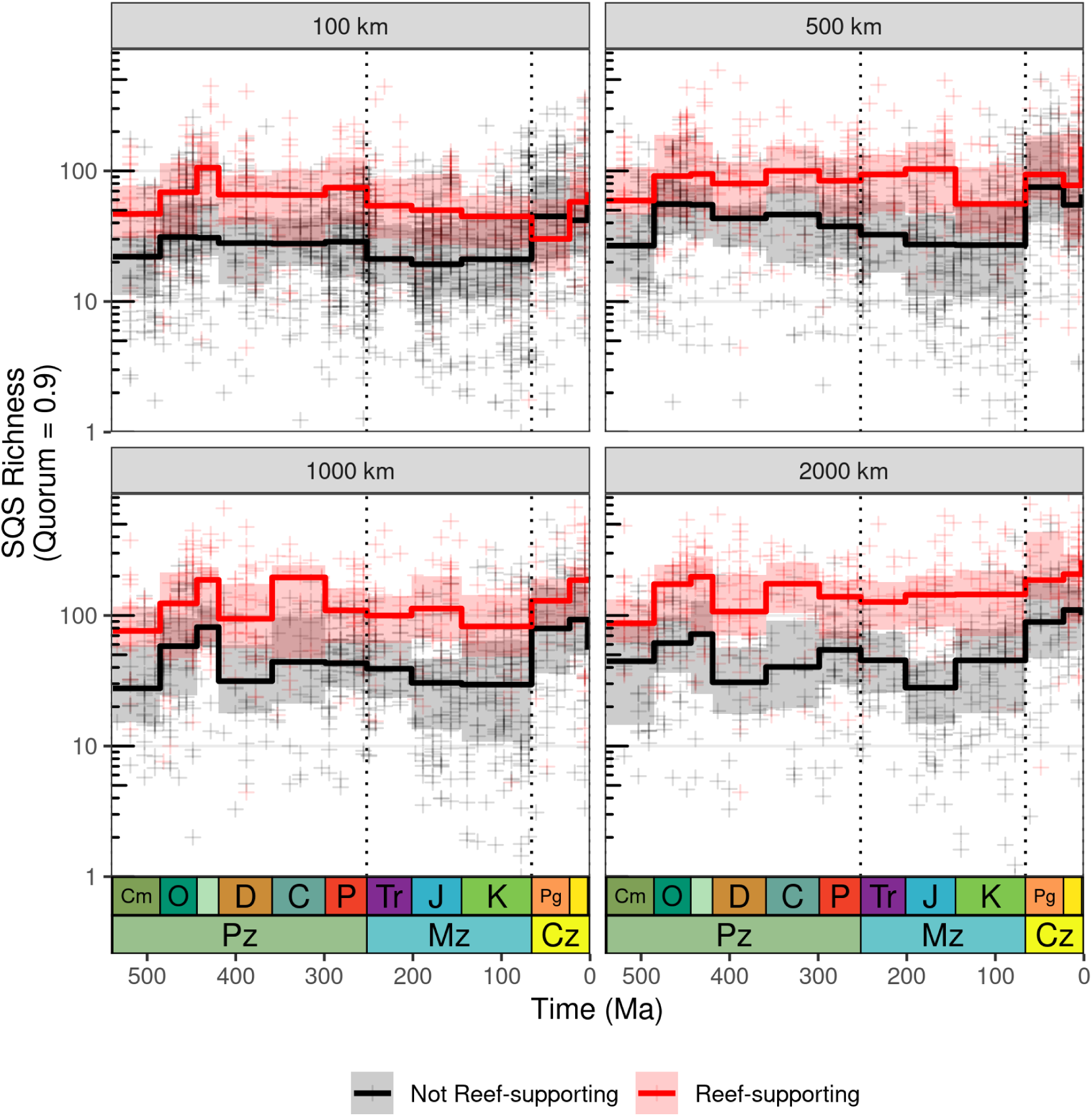
Patterns among reef-supporting and non-reef-supporting regions for all grid-cell sizes (spacings of 100 km, 500 km, 1000 km and 2000 km). Crosses represent SQS diversity estimates for individual grid cell regions, while trend lines represent medians and interquartile ranges of regional diversity for geological periods.

**Fig. S6:**
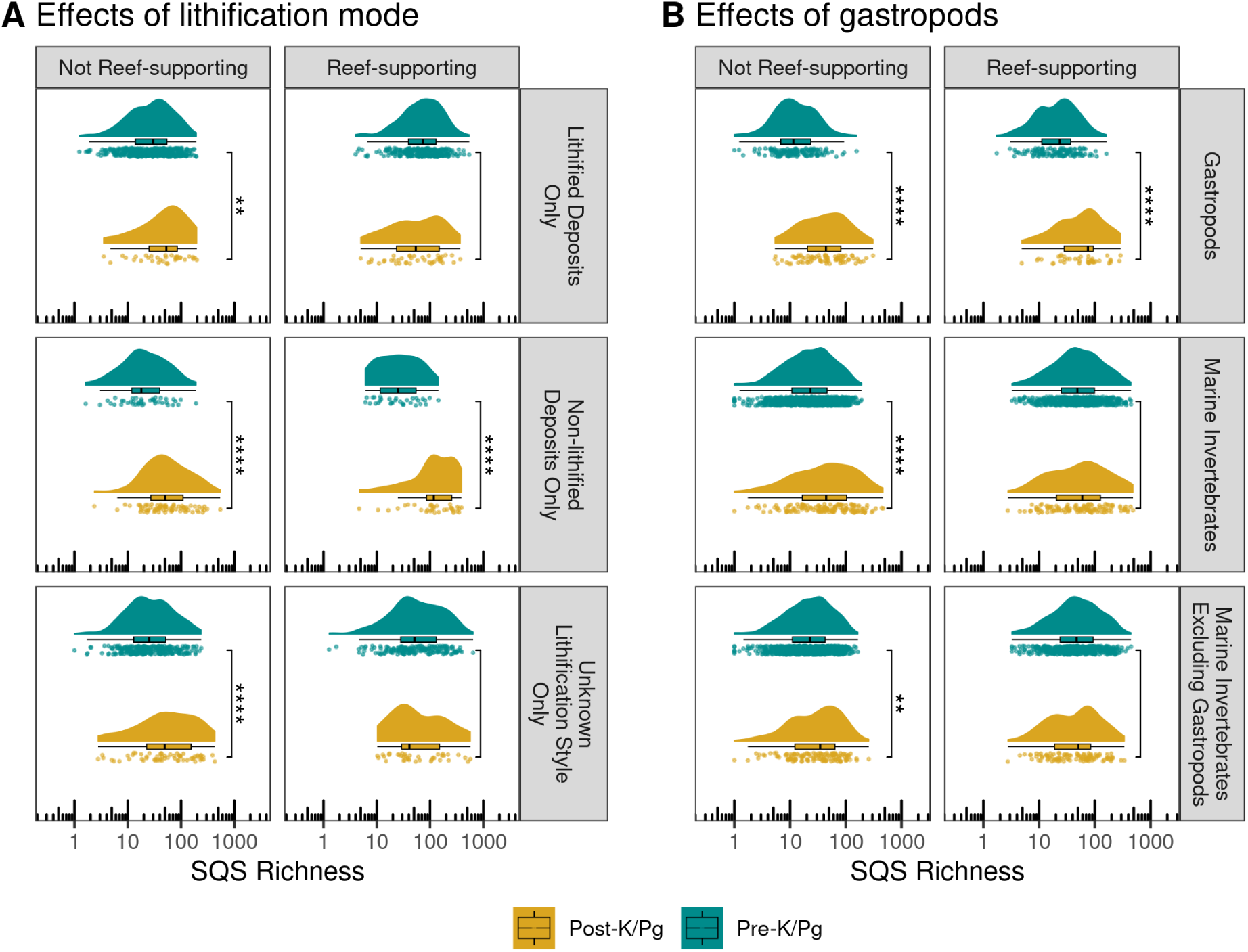
Potential drivers of the apparent increase in diversity of non-reef-supporting regions that occurred at or just before the K/Pg. (A) Effects of lithification style on diversity within reef-supporting and non-reef-supporting regions across the K/Pg boundary. (B) Effects of including or excluding gastropods on diversity in marine animals within reef-supporting and non-reef-supporting regions across the K/Pg boundary. Gastropods experienced a large increase in diversity in the latest Cretaceous, especially in non-reef-supporting regions. When gastropods are excluded from diversity estimates for marine invertebrates, little increase across the K/Pg is evident, suggesting that this event is primarily driven by the diversity dynamics of gastropods. Each plot shows three ways of visualizing the data: individual data points with jitter, a boxplot (line denotes the median value, the hinges of the box correspond to the interquartile range, and whiskers extend from the hinges to the smallest/largest values at most 1.5 * IQR of the hinge), and a density plot using a Gaussian kernel with a smoothing bandwidth of 1. Statistical significance for Wilcoxon tests between groups is indicated by either no text (p ¿ 0.05), * (p ≤ 0.05), ** (p ≤ 0.01), *** (p ≤ 0.001) or **** (p ≤ 0.0001).

**Fig. S7:**
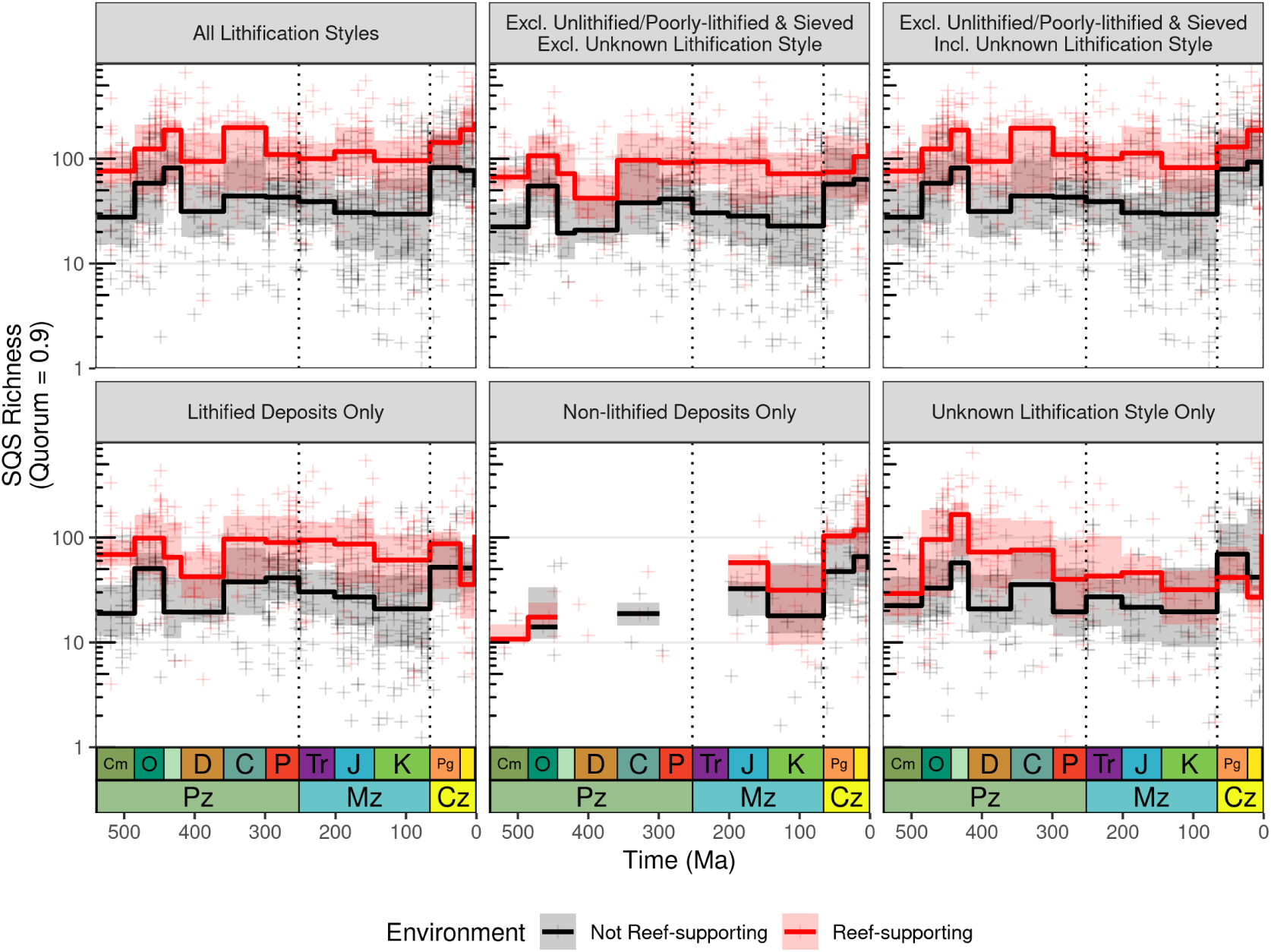
Differences in Phanerozoic marine invertebrate animal diversity patterns between reef-supporting (red) and non-reef-supporting (black) regions (equal-area hexagonal/pentagonal grid cells with spacings of 1000 km for all panels in this figure), comparing the effects of sifting criteria that vary the inclusion or exclusion of data associated with different lithification styles. See Supplementary Methods for discussion of these sifting criteria. Crosses represent SQS diversity estimates for individual grid cell regions, while trend lines represent medians and interquartile ranges of regional diversity for geological periods.

**Fig. S8:**
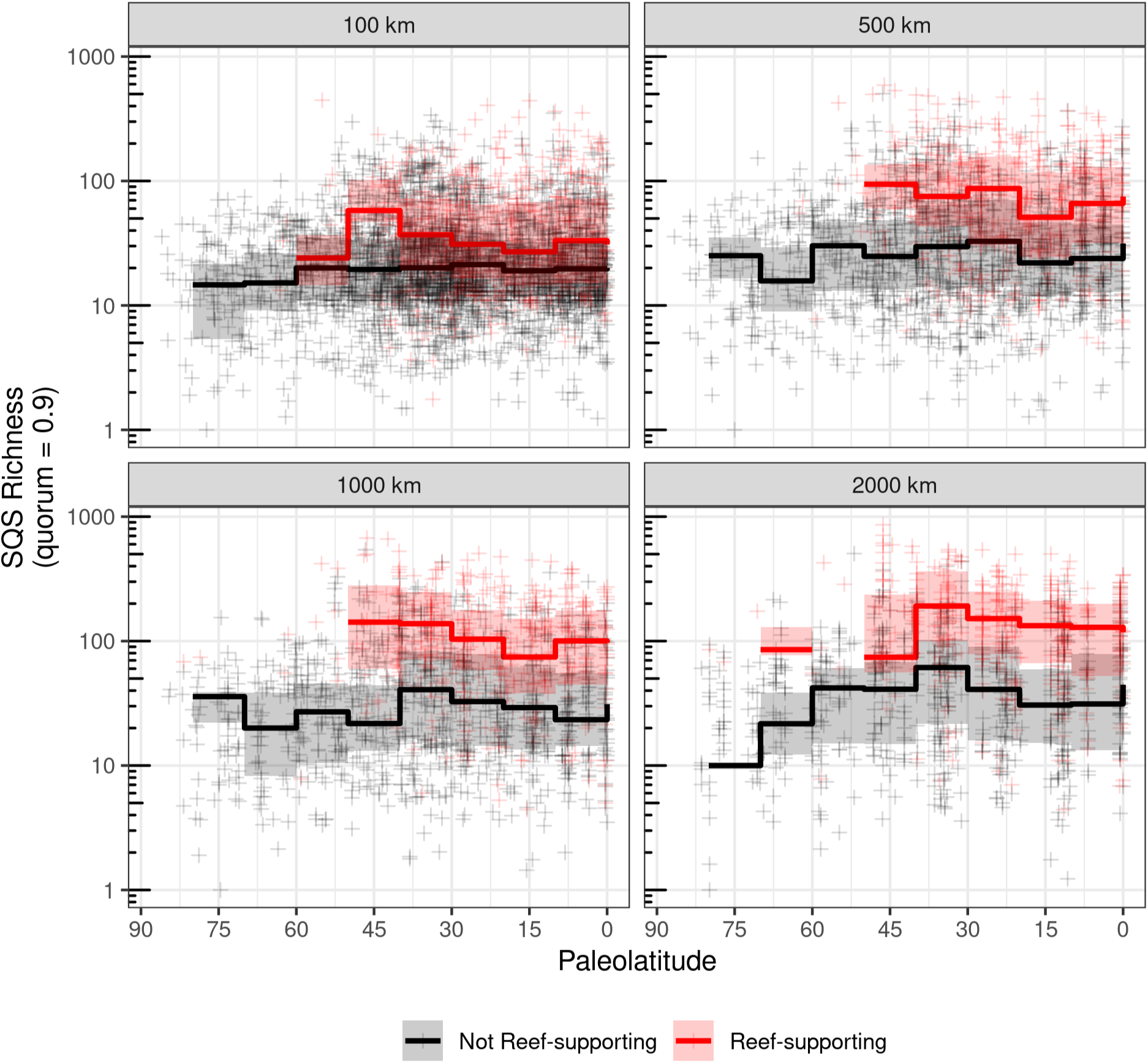
Latitudinal gradients of Phanerozoic marine invertebrate animal diversity, separately showing trends across paleolatitudes for reef-supporting (red) and non-reef-supporting (black) regions (equal-area hexagonal/pentagonal grid cells with spacings of 100 km, 500 km, 1000 km and 2000 km). Collections explicitly identified as representing unlithified or poorly-lithified-and-sieved deposits were excluded. Note logarithmic y-axis. Crosses represent SQS diversity estimates for individual grid cell regions, while trend lines represent medians and interquartile ranges of regional diversity for 10 degree paleolatitudinal bins. Diversity is significantly higher in reef-supporting regions than in non-reef-supporting regions, even at low paleolatitudes, indicating that this effect is not solely driven by higher diversity in all tropical low-paleolatitude regions.

**Fig. S9:**
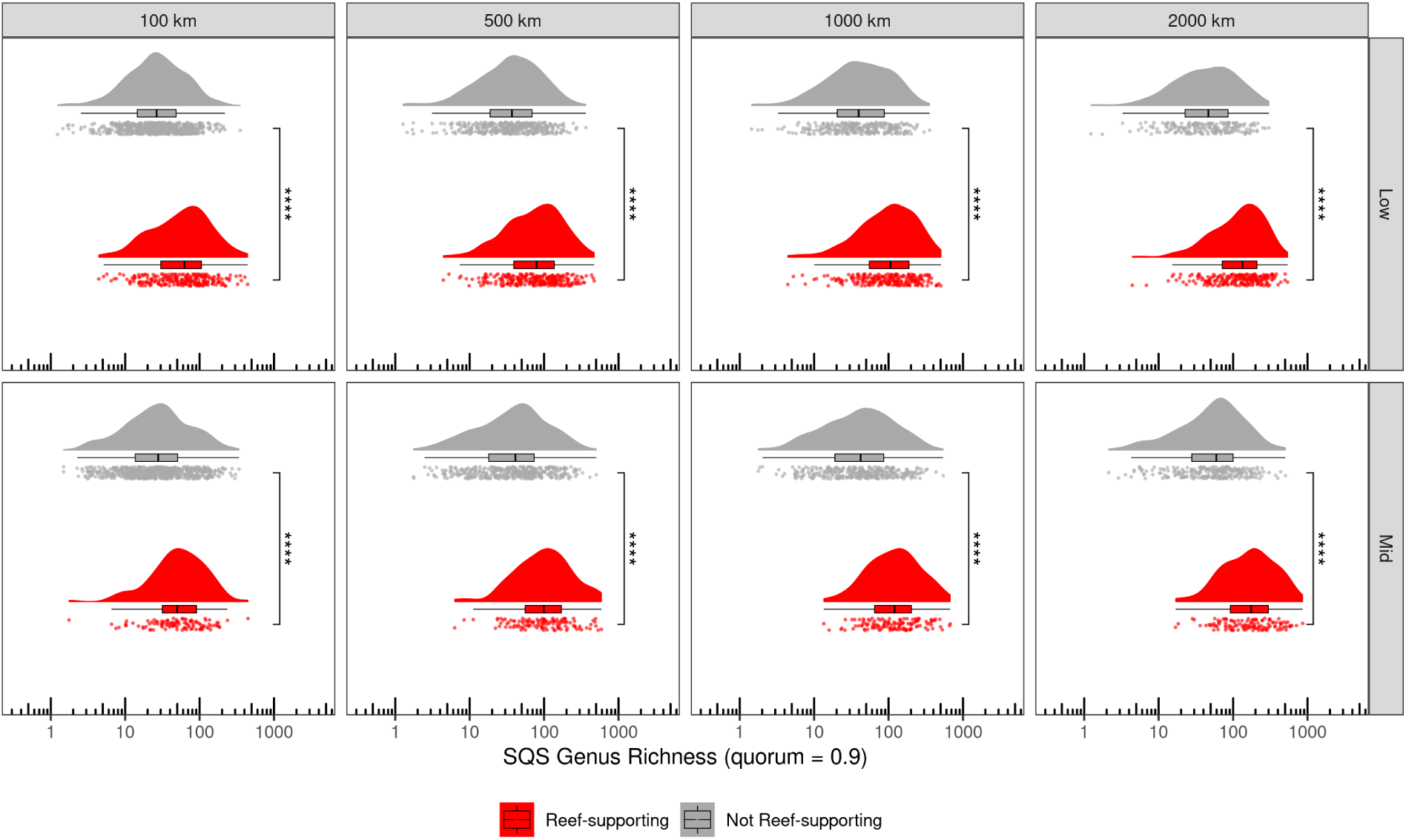
Distributions of SQS diversity (quorum = 0.9) within reef-supporting (red) and non-reef-supporting (gray) regions (equal-area hexagonal/pentagonal grid cells with 100 km, 500 km, 1000 km and 2000 km spacings) between low (0–30°) and mid (30–60°) absolute paleolatitude zones. Diversity is consistently significantly higher in reef-supporting regions than in non-reef-supporting regions, regardless of paleolatitude zones. Each plot shows three ways of visualizing the data: individual data points with jitter, a boxplot (line denotes the median value, the hinges of the box correspond to the interquartile range, and whiskers extend from the hinges to the smallest/largest values at most 1.5 * IQR of the hinge), and a density plot using a Gaussian kernel with a smoothing bandwidth of 1. Statistical significance for Wilcoxon tests between groups is indicated by either no text (p ¿ 0.05), * (p ≤ 0.05), ** (p ≤ 0.01), *** (p ≤ 0.001) or **** (p ≤ 0.0001).

**Fig. S10:**
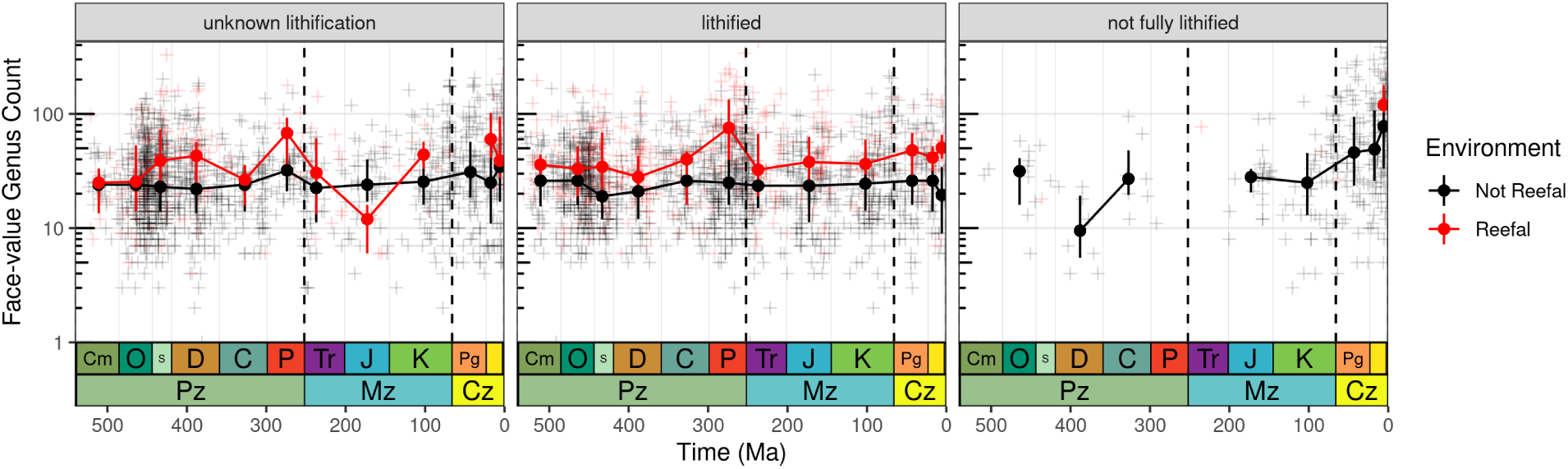
Per-formation counts of genera for reef-supporting and non-reef-supporting geological formations, broken down by lithification style. To simplify the presentation of the data, panels show data of unknown lithification only, lithified data only, and data that is not fully lithified only, including unlithified and poorly-lithified deposits. Crosses represent counts of genera for individual formations, while trend-lines indicate medians and interquartile ranges for geological periods. Quality criteria were used to exclude poorly-sampled formations (at least 20 collections, five references or four higher taxa).

**Fig. S11:**
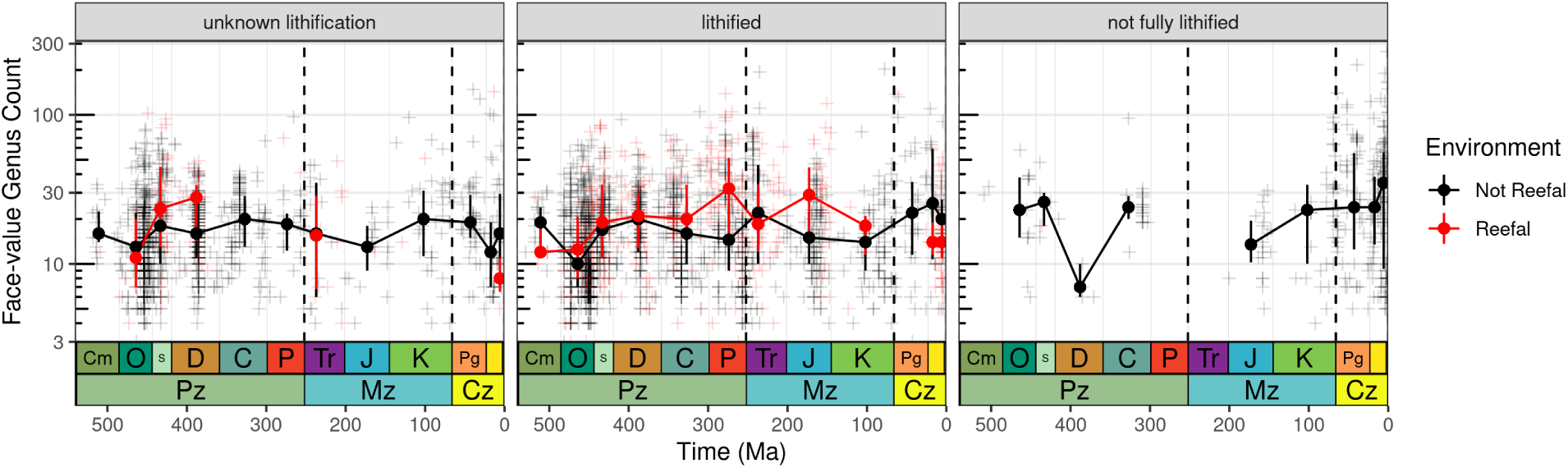
Per-collection counts of genera for reefal and non-reefal facies, broken down by lithification style. To simplify the presentation of the data, panels show data of unknown lithification only, lithified data only, and data that is not fully lithified only, including unlithified and poorly-lithified deposits. Crosses represent counts of genera for individual collections, while trend-lines indicate medians and interquartile ranges for geological periods. Quality criteria were used to exclude poorly-sampled collections (at least five references per collection or four higher taxa).

**Fig. S12:**
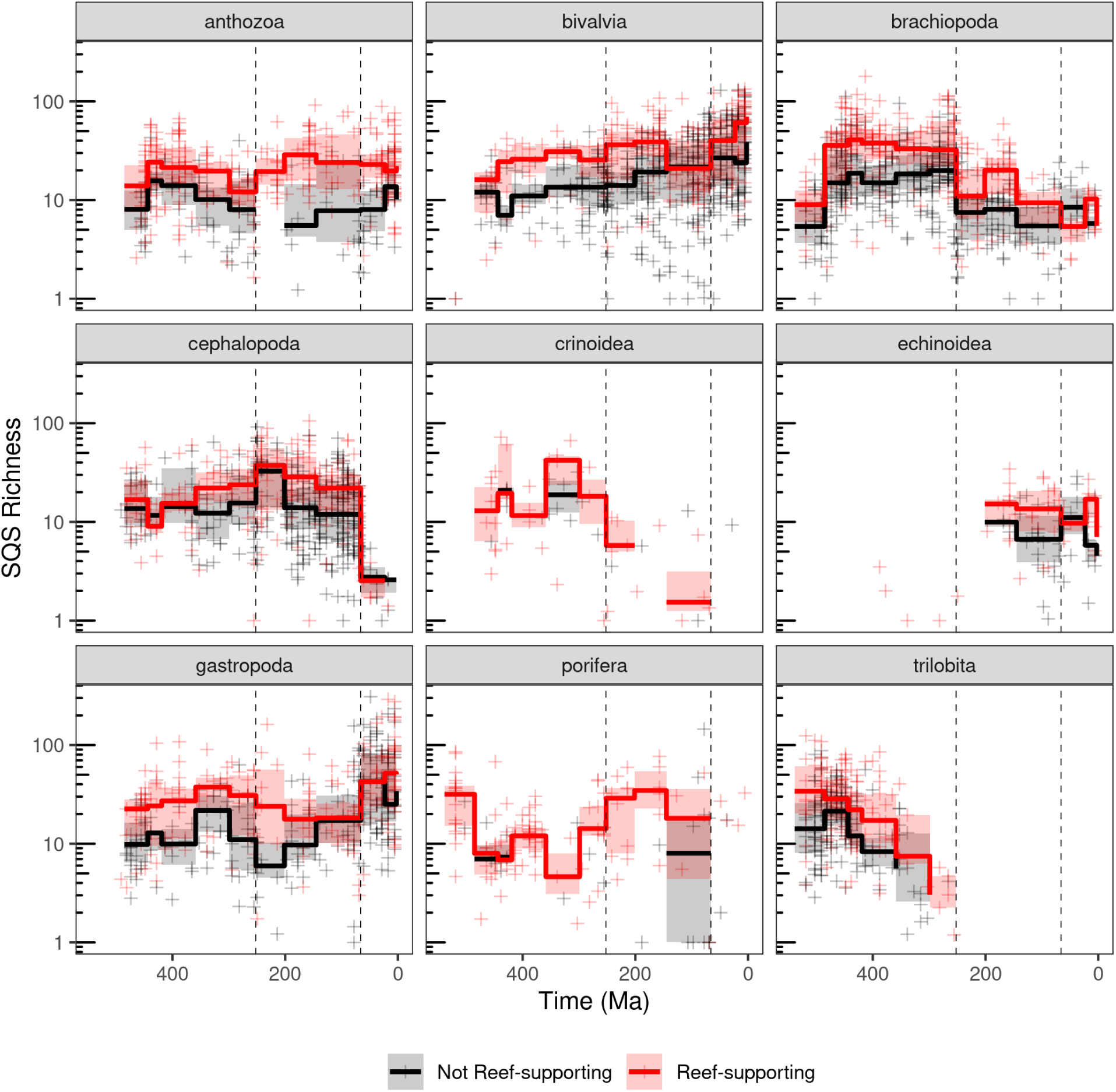
Differences in diversity patterns for major clades of marine invertebrate animals between reef-supporting (red) and non-reef-supporting (black) regions (equal-area hexagonal/pentagonal grid cells with 1000 km spacings). Note logarithmic y-axes. Crosses represent SQS diversity estimates for individual grid cell regions, while trend lines represent medians and interquartile ranges of regional diversity for geological periods. Dashed lines represent boundaries between geological eras. PaleoDB collection data excludes those identified as unlithified and poorly-lithified-and-sieved deposits, but includes collections with no information on lithification style.

**Fig. S13:**
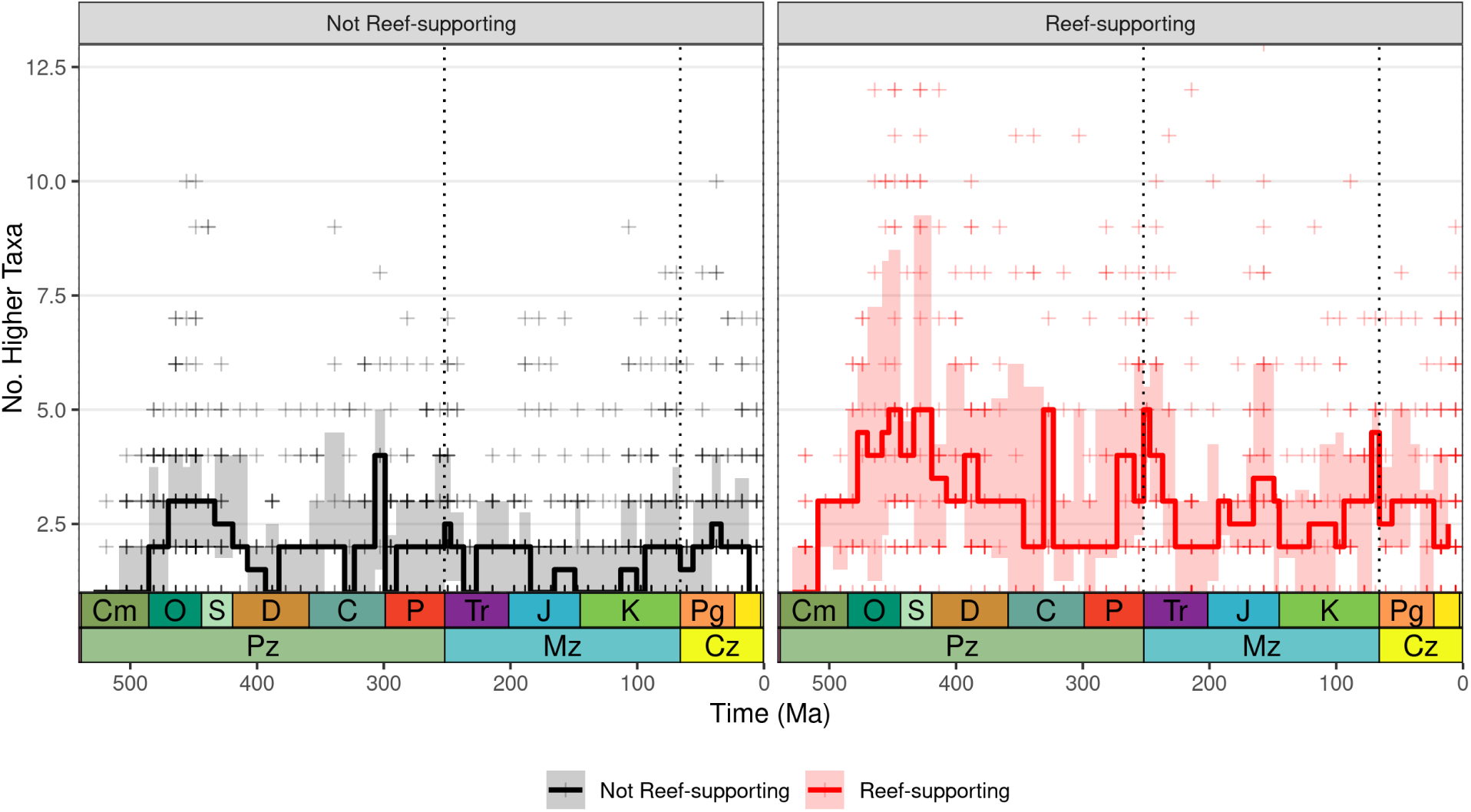
Counts of higher taxa (major groups of taxa comprising bivalvia, rhynchonelliformea, linguliformea, cephalopoda, gastropoda, chordata sans tetrapoda, anthozoa, trilobita, conodonta, bryozoa, porifera, tetrapoda, crinoidea, echinoidea, graptolithina, decapoda, annelida) for reef-supporting (red) and non-reef-supporting (black) regions (1000 km equal-area hexagonal grid cells). Crosses represent face-value counts of these higher taxa for grid cells, and lines with transparent ribbons represent medians and interquartile ranges for geological periods.

**Fig. S14:**
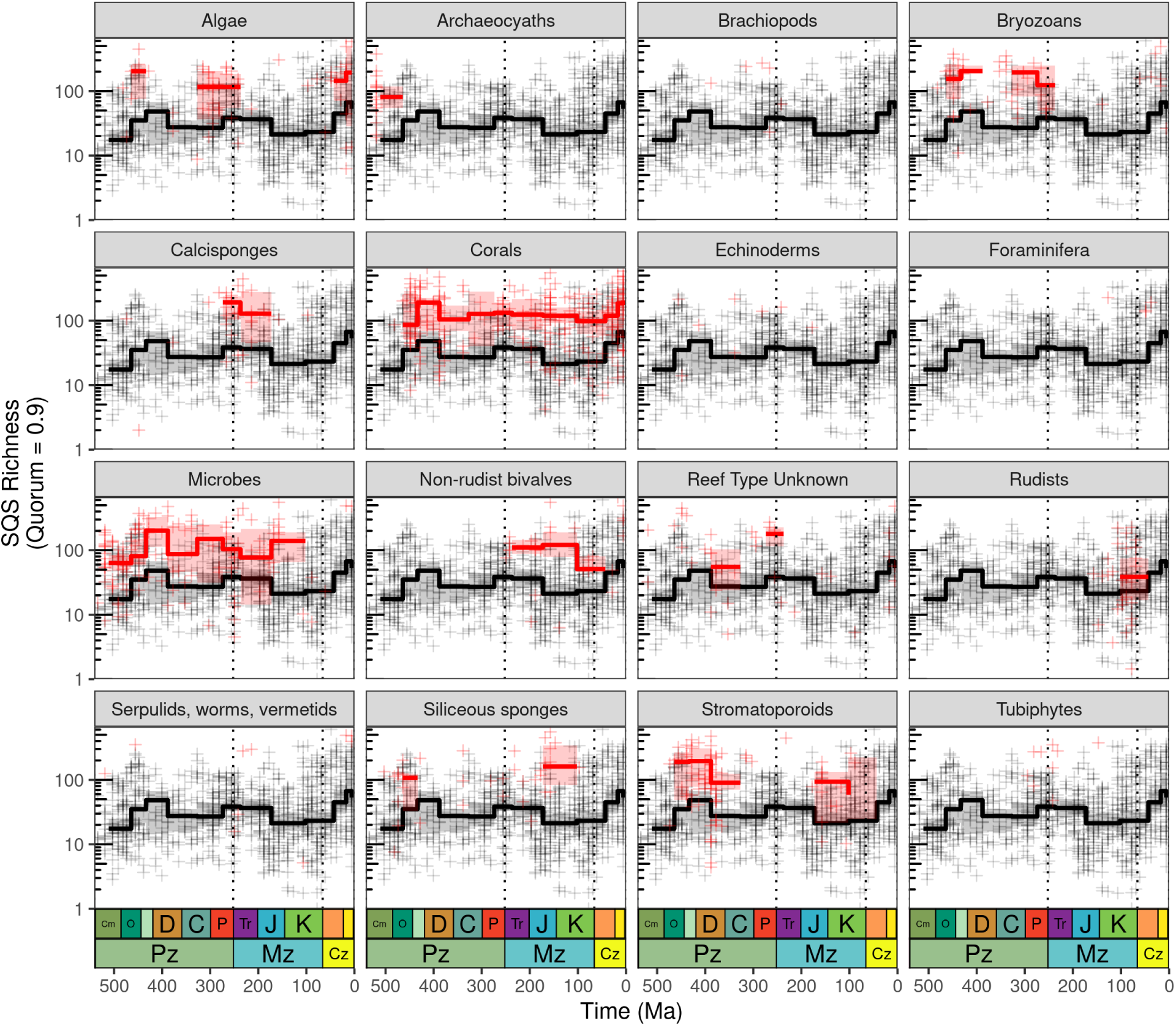
Differences in diversity patterns (SQS, quorum = 0.9) for marine invertebrate animals between reef-supporting (red) and non-reef-supporting (black) regions (equal-area hexagonal/pentagonal grid cells with 1000 km spacings), showing patterns for additional kinds of reef-building organisms. Reef-supporting cells were identified as being associated with each particular kind of reef-building organism using the palaeocoordinates of reef sites listed in the PARED PaleoReefs database (*36*) (grid cells can appear in more than one panel if they contain more than one type of reef-building organism). Note logarithmic y-axes. Crosses represent SQS diversity estimates for individual grid cell regions, while trend lines represent medians and interquartile ranges of regional diversity for geological periods. Dashed lines represent boundaries between geological eras. PaleoDB collection data excludes those identified as unlithified and poorly-lithified-and-sieved deposits, but includes collections with no information on lithification style.

**Fig. S15:**
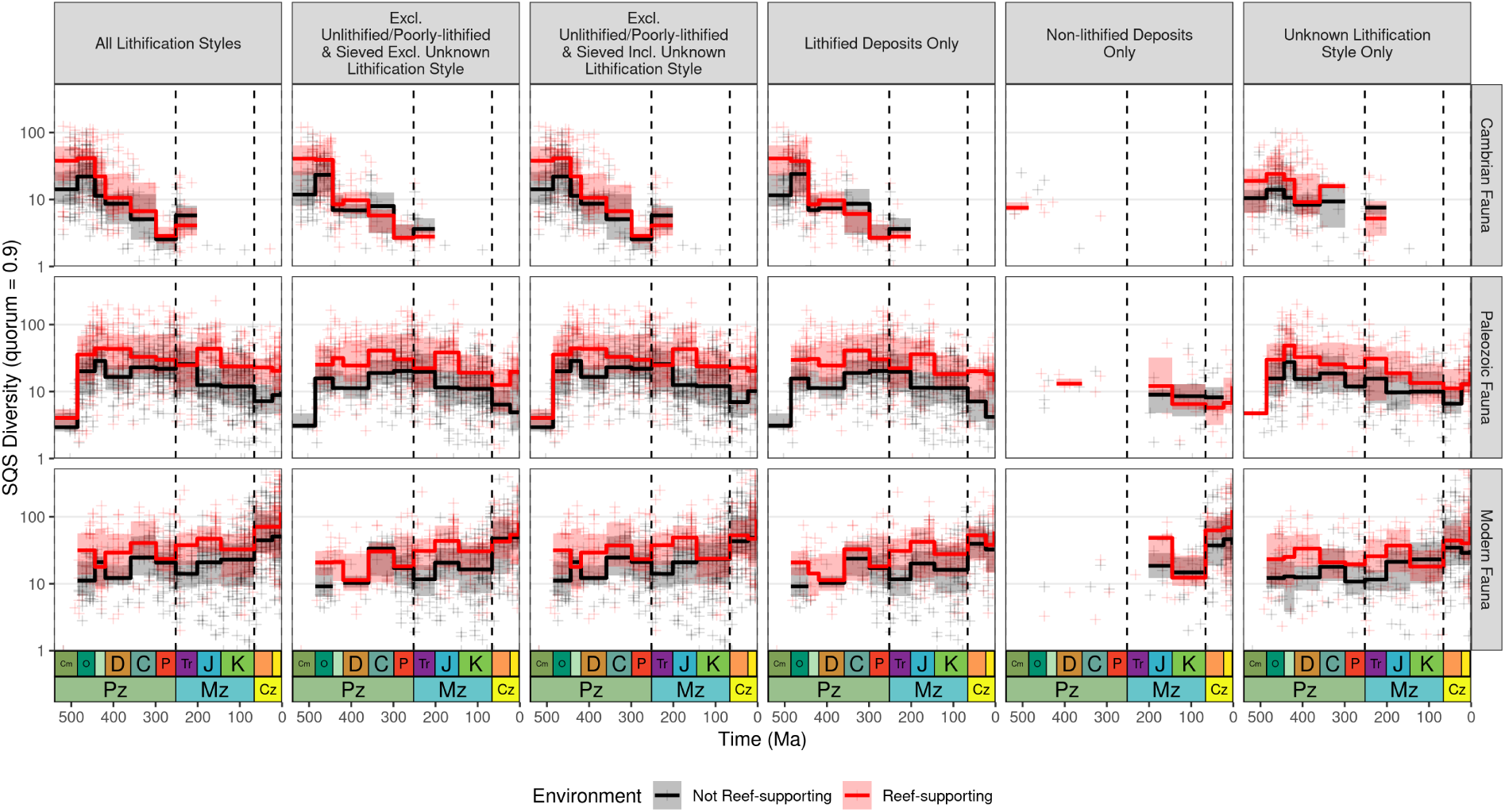
SQS diversity for reef-supporting (red) and non-reef-supporting (black) regions (equal-area hexagonal/pentagonal grid cells with 1000 km spacings) within Sepkoski’s three evolutionary faunas, partitioned by lithification style into PaleoDB collections representing lithified deposits, or those representing poorly-lithified or unlithified deposits.

**Fig. S16:**
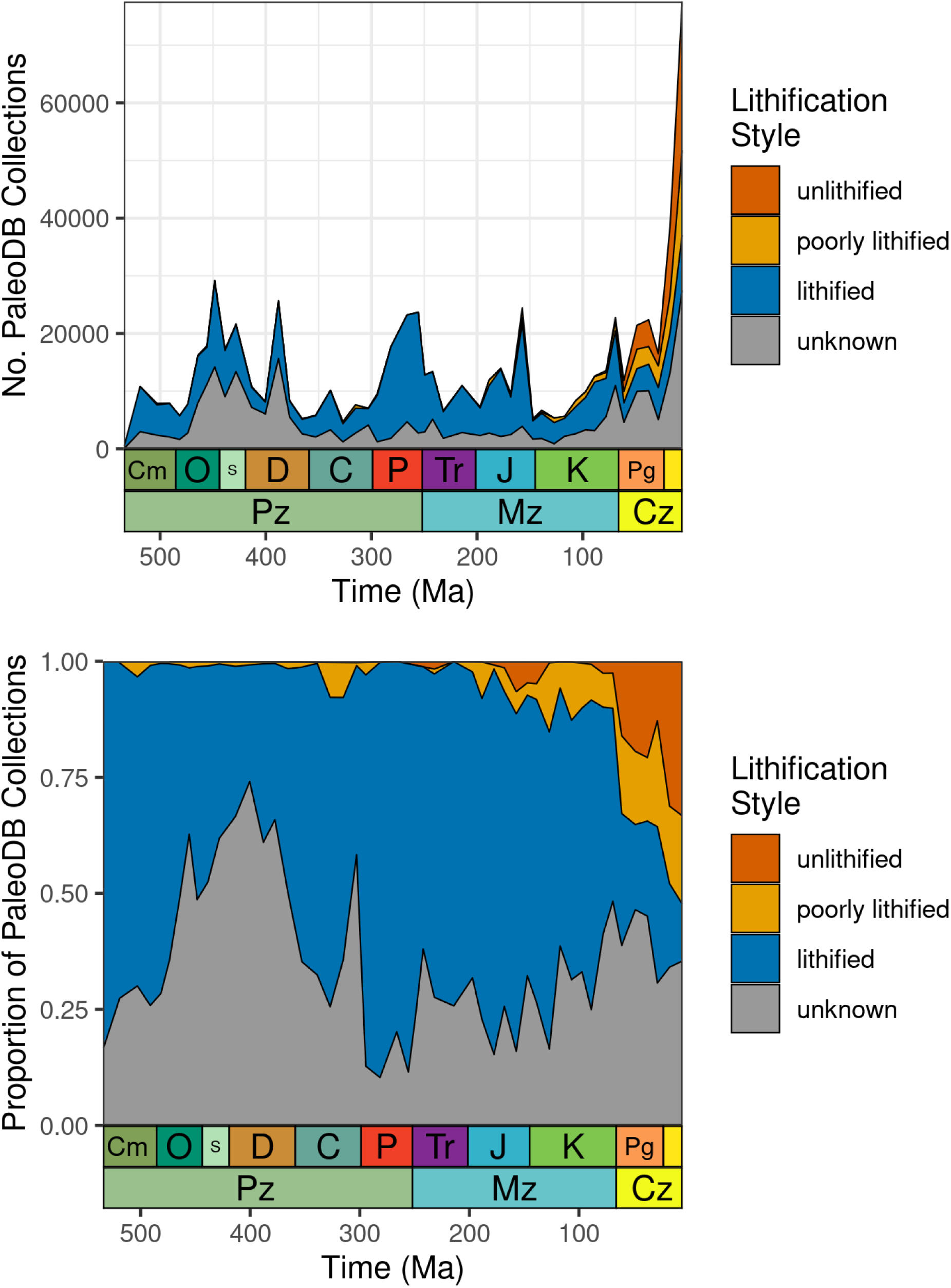
Trends in lithification style through the Phanerozoic, showing absolute counts of 100 km equal-area hexagonal/pentagonal grid cells containing fossil collections ascribed to each lithification style (unlithified, poorly-lithified, or lithified), or proportions of these counts.

**Fig. S17:**
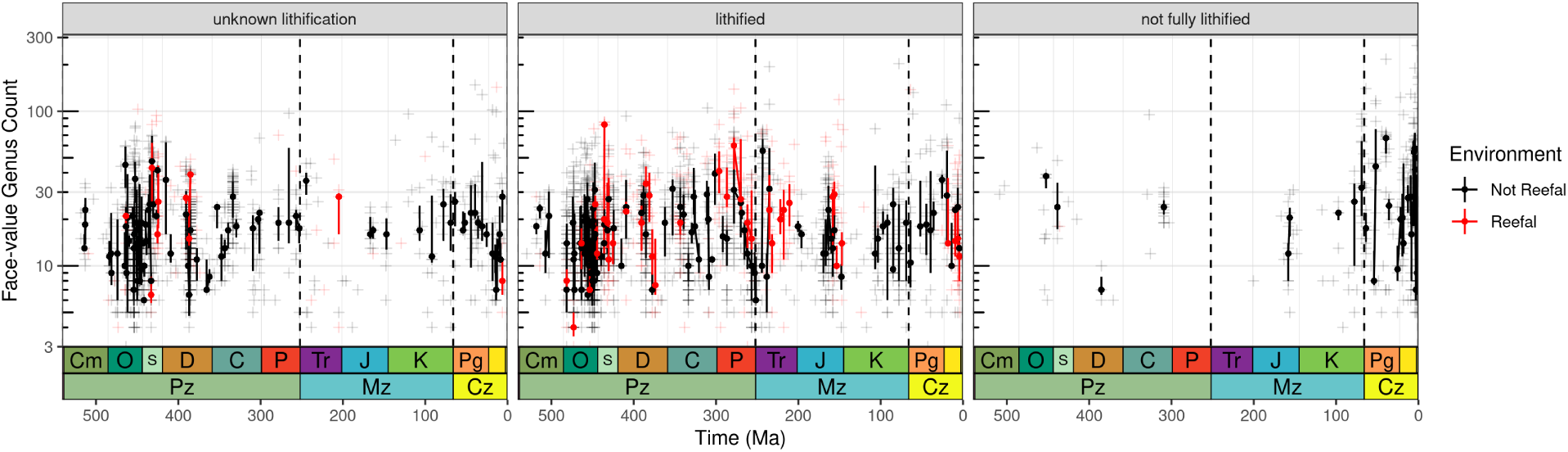
Per-collection counts of genera for reefal and non-reefal facies, broken down by lithification style. Crosses represent counts of genera for individual collections, while trend-lines indicate medians and interquartile ranges for geological stages. See Supplementary Methods for details of quality criteria used to remove uninformative collections.

**Fig. S18:**
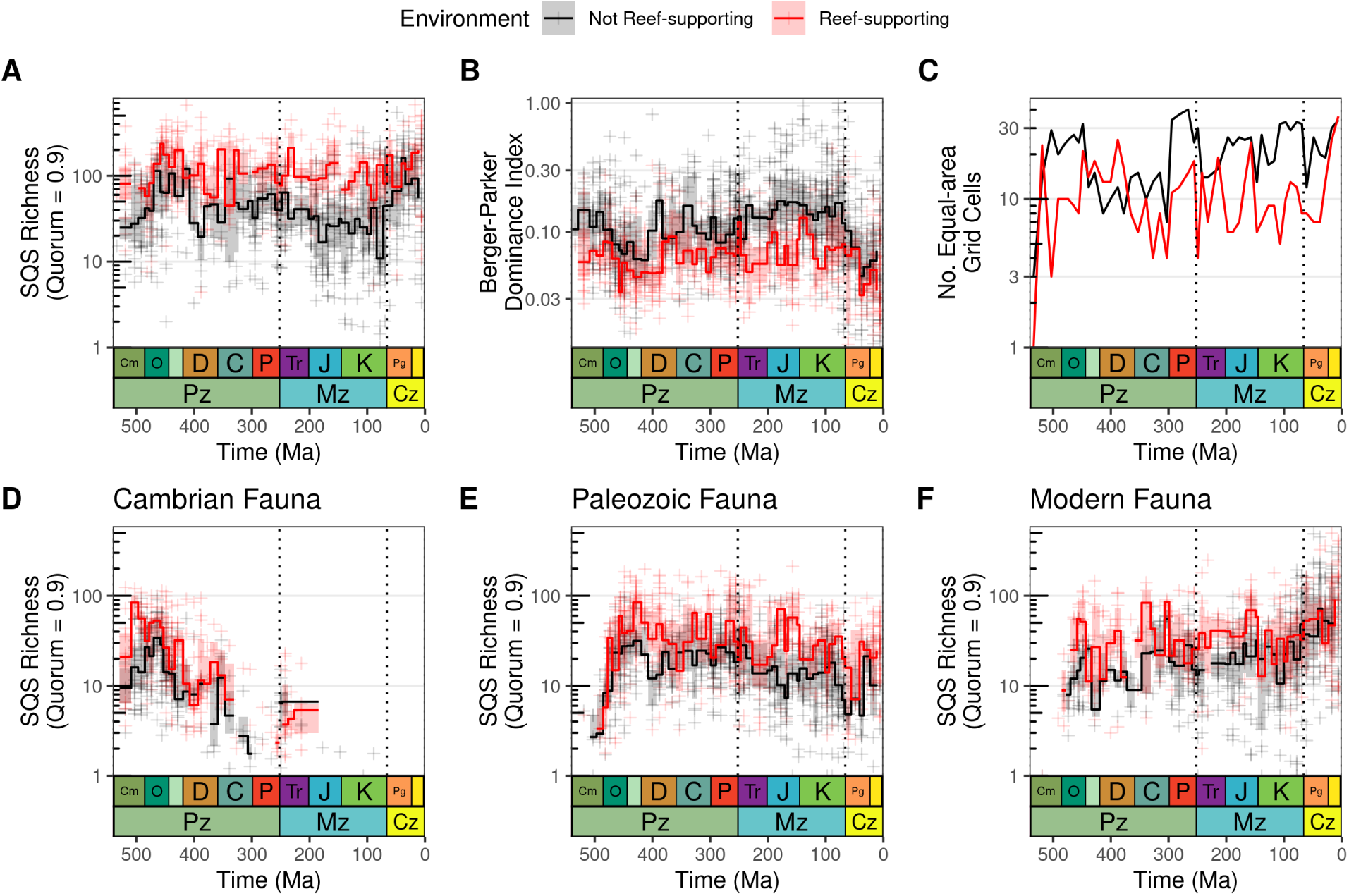
Differences in Phanerozoic marine invertebrate animal diversity patterns between reef-supporting (red) and non-reef-supporting (black) regions (equal-area hexagonal/pentagonal grid cells with spacings of 1000 km for all panels in this figure), excluding collections explicitly identified as representing unlithified or poorly-lithified-and-sieved deposits, but retaining collections that have no information about lithification style. Dotted lines represent boundaries between geological eras. Note logarithmic y-axes. For panels A–B and D–F, crosses represent SQS diversity estimates for individual grid cell regions, while trend lines represent medians and interquartile ranges of regional diversity for equal-length bins. (A) Spatially-standardized Phanerozoic marine animal diversity, contrasting patterns for reef-supporting and non-reef-supporting regions. Note that in reef-supporting regions, levels of diversity have been broadly similar since the Ordovician, with no evidence for long-term, secular trends. In non-reef-supporting regions, by contrast, levels of diversity were similar from the Ordovician to the latest Cretaceous, when diversity rose fairly rapidly to a new, higher level that was sustained through the Cenozoic. However, this K/Pg increase is strongly associated with gastropods and unlithified sediments (see fig. S6). (B) Evenness, estimated using Berger-Parker dominance index (*35*), in reef-supporting and non-reef-supporting grid cells. (C) Counts of reef-supporting and non-reef-supporting cells through the Phanerozoic, using equal-length time bins. Panels (D–F) show patterns for Sepkoski’s evolutionary faunas. (D) Cambrian Fauna (Trilobita, Linguliformea, Graptolithina, Conodonta); (E) Modern Fauna (Anthozoa, Ostracoda, Rhynchonelliformea, Cephalopoda, Crinoidea); (F) Modern Fauna (Bryozoa, Bivalvia, Gastropoda, Echinoidea, Chondrichthyes).

**Fig. S19:**
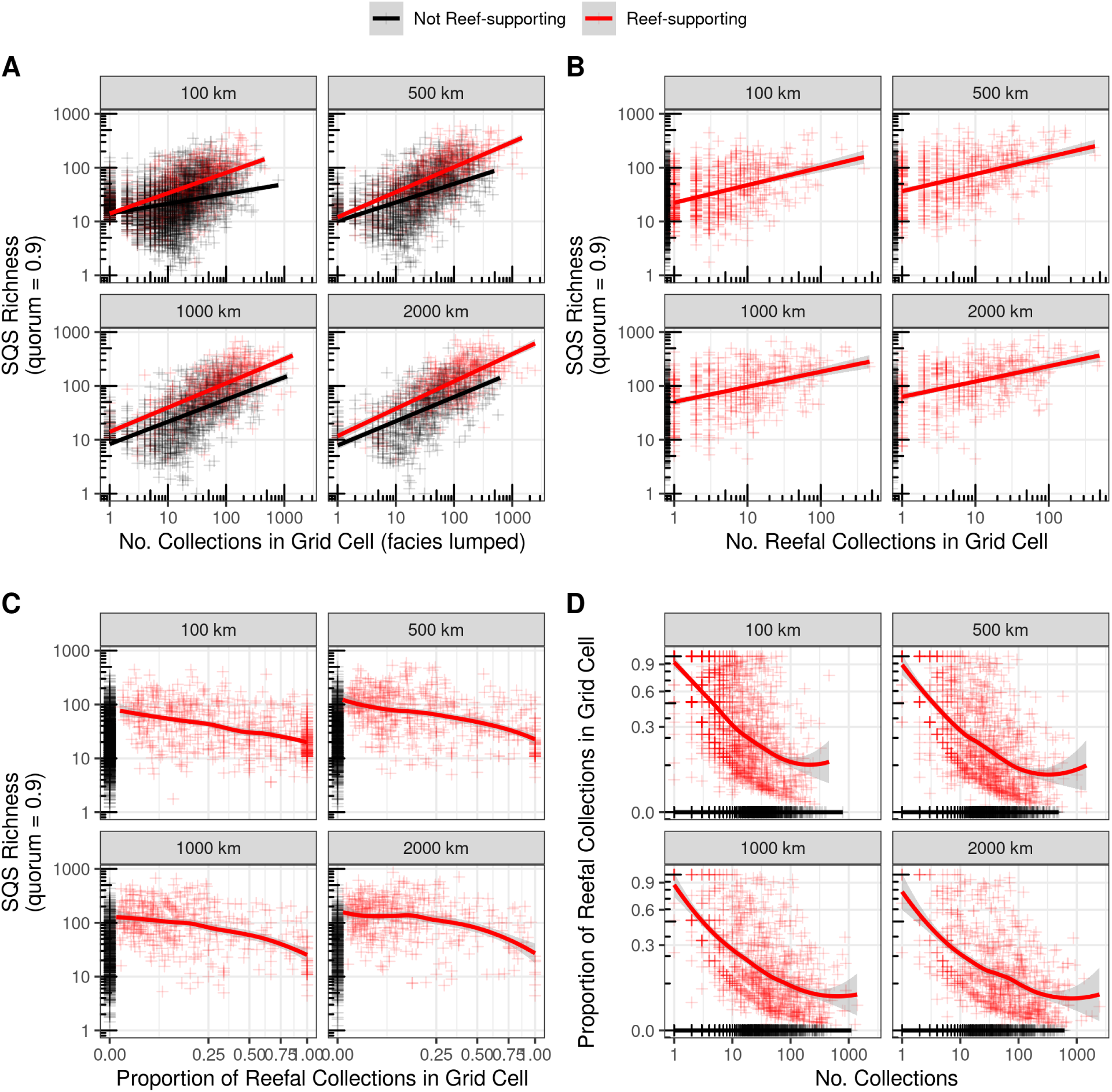
**Relationships between collection counts, SQS diversity, and proportions of reefal collections.** Relationships between (A) diversity (SQS, quorum = 0.9) and the total number of collections of any facies present in a region; (B) diversity (SQS, quorum = 0.9) and the number of reefal-type collections present in a region; (C) diversity (SQS, quorum = 0.9) and the proportion of reefal-type collections (versus non-reefal) present in a region; and (D) the proportion of reefal-type collections (versus non-reefal) and counts of collections present in a region, all for reef-supporting (red) and non-reef-supporting (black) regions. Regions represent equal-area hexagonal/pentagonal grid cells with 100 km, 500 km, 1000 km and 2000 km spacings. Lines represent OLS linear model fits in panels A and B and LOESS fits in panels C and D. Counts of collections and SQS diversity have log-transformed axes, while proportions of reefal collections are square-root–transformed. Crosses represent SQS diversity estimates for individual grid cell regions. PaleoDB collection data excludes those identified as unlithified and poorly-lithified-and-sieved deposits, but includes collections with no information on lithification style. Non-reef-supporting data points are confined to the left-hand side of the plot in panels B and C, and to the bottom of the plot in panel D, since they lack any reefal collections. Panel (C) shows that diversity declines as the proportion of reefal collections increases. This is because regions containing a high proportion of reefal collections also tend to have a smaller absolute number of collections, as shown in panel D.

**Table S1:** ANOVA with Tukey HSD for collection-subsampled reefal/non-reefal counts of genera from reef-supporting/not cells.

| contrast | null.value | estimate | conf.low | conf.high | adj.p.value | significance |
| --- | --- | --- | --- | --- | --- | --- |
| **100km cells, 10 collection quota** |  |  |  |  |  |  |
| Non-reefal Facies in Reef-supporting Regions — Non-reefal Facies in Non-reef-supporting Regions | 0 | 0.2264219 | 0.1000974 | 0.3527463 | 0.0000853 | ¡ 0.05 |
| Reefal Facies in Reef-supporting Regions — Non-reefal Facies in Non-reef-supporting Regions | 0 | 0.4826773 | 0.2936908 | 0.6716638 | 0.0000000 | ¡ 0.05 |
| Reefal Facies in Reef-supporting Regions — Non-reefal Facies in Reef-supporting Regions | 0 | 0.2562554 | 0.0528080 | 0.4597029 | 0.0089573 | ¡ 0.05 |
| **100km cells, 20 collection quota** |  |  |  |  |  |  |
| Non-reefal Facies in Reef-supporting Regions — Non-reefal Facies in Non-reef-supporting Regions | 0 | 0.3700756 | 0.2260246 | 0.5141265 | 0.0000000 | ¡ 0.05 |
| Reefal Facies in Reef-supporting Regions — Non-reefal Facies in Non-reef-supporting Regions | 0 | 0.5598309 | 0.3177896 | 0.8018721 | 0.0000003 | ¡ 0.05 |
| Reefal Facies in Reef-supporting Regions — Non-reefal Facies in Reef-supporting Regions | 0 | 0.1897553 | -0.0665852 | 0.4460958 | 0.1912034 | n.s. |
| **100km cells, 40 collection quota** |  |  |  |  |  |  |
| Non-reefal Facies in Reef-supporting Regions — Non-reefal Facies in Non-reef-supporting Regions | 0 | 0.4182346 | 0.2353679 | 0.6011014 | 0.0000006 | ¡ 0.05 |
| Reefal Facies in Reef-supporting Regions — Non-reefal Facies in Non-reef-supporting Regions | 0 | 0.4966521 | 0.1447910 | 0.8485131 | 0.0029523 | ¡ 0.05 |
| Reefal Facies in Reef-supporting Regions — Non-reefal Facies in Reef-supporting Regions | 0 | 0.0784174 | -0.2861249 | 0.4429598 | 0.8673890 | n.s. |
| **500km cells, 10 collection quota** |  |  |  |  |  |  |
| Non-reefal Facies in Reef-supporting Regions — Non-reefal Facies in Non-reef-supporting Regions | 0 | 0.1517835 | 0.0456124 | 0.2579545 | 0.0023711 | ¡ 0.05 |
| Reefal Facies in Reef-supporting Regions — Non-reefal Facies in Non-reef-supporting Regions | 0 | 0.4143076 | 0.2668829 | 0.5617323 | 0.0000000 | ¡ 0.05 |
| Reefal Facies in Reef-supporting Regions — Non-reefal Facies in Reef-supporting Regions | 0 | 0.2625242 | 0.1093486 | 0.4156997 | 0.0001834 | ¡ 0.05 |
| **500km cells, 20 collection quota** |  |  |  |  |  |  |
| Non-reefal Facies in Reef-supporting Regions — Non-reefal Facies in Non-reef-supporting Regions | 0 | 0.2202123 | 0.1042036 | 0.3362210 | 0.0000289 | ¡ 0.05 |
| Reefal Facies in Reef-supporting Regions — Non-reefal Facies in Non-reef-supporting Regions | 0 | 0.3879187 | 0.2036381 | 0.5721993 | 0.0000029 | ¡ 0.05 |
| Reefal Facies in Reef-supporting Regions — Non-reefal Facies in Reef-supporting Regions | 0 | 0.1677064 | -0.0196028 | 0.3550157 | 0.0899457 | n.s. |
| **500km cells, 40 collection quota** |  |  |  |  |  |  |
| Non-reefal Facies in Reef-supporting Regions — Non-reefal Facies in Non-reef-supporting Regions | 0 | 0.3303667 | 0.1887091 | 0.4720243 | 0.0000002 | ¡ 0.05 |
| Reefal Facies in Reef-supporting Regions — Non-reefal Facies in Non-reef-supporting Regions | 0 | 0.5517774 | 0.2838576 | 0.8196972 | 0.0000057 | ¡ 0.05 |
| Reefal Facies in Reef-supporting Regions — Non-reefal Facies in Reef-supporting Regions | 0 | 0.2214107 | -0.0473561 | 0.4901775 | 0.1293051 | n.s. |
| **1000km cells, 10 collection quota** |  |  |  |  |  |  |
| Non-reefal Facies in Reef-supporting Regions — Non-reefal Facies in Non-reef-supporting Regions | 0 | 0.0953068 | -0.0116248 | 0.2022384 | 0.0919579 | n.s. |
| Reefal Facies in Reef-supporting Regions — Non-reefal Facies in Non-reef-supporting Regions | 0 | 0.3441853 | 0.1990279 | 0.4893427 | 0.0000001 | ¡ 0.05 |
| Reefal Facies in Reef-supporting Regions — Non-reefal Facies in Reef-supporting Regions | 0 | 0.2488785 | 0.1010565 | 0.3967004 | 0.0002461 | ¡ 0.05 |
| **1000km cells, 20 collection quota** |  |  |  |  |  |  |
| Non-reefal Facies in Reef-supporting Regions — Non-reefal Facies in Non-reef-supporting Regions | 0 | 0.1434669 | 0.0240526 | 0.2628811 | 0.0136115 | ¡ 0.05 |
| Reefal Facies in Reef-supporting Regions — Non-reefal Facies in Non-reef-supporting Regions | 0 | 0.3251909 | 0.1406063 | 0.5097756 | 0.0001174 | ¡ 0.05 |
| Reefal Facies in Reef-supporting Regions — Non-reefal Facies in Reef-supporting Regions | 0 | 0.1817241 | -0.0022354 | 0.3656835 | 0.0536965 | n.s. |
| **1000km cells, 40 collection quota** |  |  |  |  |  |  |
| Non-reefal Facies in Reef-supporting Regions — Non-reefal Facies in Non-reef-supporting Regions | 0 | 0.2820043 | 0.1417831 | 0.4222254 | 0.0000093 | ¡ 0.05 |
| Reefal Facies in Reef-supporting Regions — Non-reefal Facies in Non-reef-supporting Regions | 0 | 0.4751477 | 0.2310287 | 0.7192668 | 0.0000187 | ¡ 0.05 |
| Reefal Facies in Reef-supporting Regions — Non-reefal Facies in Reef-supporting Regions | 0 | 0.1931435 | -0.0477188 | 0.4340057 | 0.1439121 | n.s. |
| **2000km cells, 10 collection quota** |  |  |  |  |  |  |
| Non-reefal Facies in Reef-supporting Regions — Non-reefal Facies in Non-reef-supporting Regions | 0 | 0.1411193 | 0.0386009 | 0.2436377 | 0.0036549 | ¡ 0.05 |
| Reefal Facies in Reef-supporting Regions — Non-reefal Facies in Non-reef-supporting Regions | 0 | 0.4009389 | 0.2661772 | 0.5357006 | 0.0000000 | ¡ 0.05 |
| Reefal Facies in Reef-supporting Regions — Non-reefal Facies in Reef-supporting Regions | 0 | 0.2598196 | 0.1262098 | 0.3934294 | 0.0000171 | ¡ 0.05 |
| **2000km cells, 20 collection quota** |  |  |  |  |  |  |
| Non-reefal Facies in Reef-supporting Regions — Non-reefal Facies in Non-reef-supporting Regions | 0 | 0.1628196 | 0.0505478 | 0.2750915 | 0.0020161 | ¡ 0.05 |
| Reefal Facies in Reef-supporting Regions — Non-reefal Facies in Non-reef-supporting Regions | 0 | 0.3853376 | 0.2243640 | 0.5463113 | 0.0000001 | ¡ 0.05 |
| Reefal Facies in Reef-supporting Regions — Non-reefal Facies in Reef-supporting Regions | 0 | 0.2225180 | 0.0647863 | 0.3802498 | 0.0027901 | ¡ 0.05 |
| **2000km cells, 40 collection quota** |  |  |  |  |  |  |
| Non-reefal Facies in Reef-supporting Regions — Non-reefal Facies in Non-reef-supporting Regions | 0 | 0.3000747 | 0.1601274 | 0.4400220 | 0.0000021 | ¡ 0.05 |
| Reefal Facies in Reef-supporting Regions — Non-reefal Facies in Non-reef-supporting Regions | 0 | 0.5615642 | 0.3279159 | 0.7952125 | 0.0000001 | ¡ 0.05 |
| Reefal Facies in Reef-supporting Regions — Non-reefal Facies in Reef-supporting Regions | 0 | 0.2614895 | 0.0367220 | 0.4862570 | 0.0177708 | ¡ 0.05 |

**Table S2:** Wilcoxon tests of whether there is a statistically significant difference in diversity among reef-supporting/non-reef-supporting regions, for the entire Phanerozoic, for several different sifting criteria (see Supplementary Methods) and equal-area hexagonal/pentagonal grid-cell sizes (spacings of 100 km, 500 km, 1000 km and 2000 km). Magnitude denotes strength of effect size.

| grid cell size | group1 | group2 | n1 | n2 | statistic | p | p.signif | effect size (r) | magnitude | Median SQS<br>(Not Reef-supporting) | Median SQS<br>(Reef-supporting) | Richness Ratio |
| --- | --- | --- | --- | --- | --- | --- | --- | --- | --- | --- | --- | --- |
| **All Lithification Styles** |  |  |  |  |  |  |  |  |  |  |  |  |
| 100km | Not Reef-supporting | Reef-supporting | 2211 | 588 | 159876.5 | 0.00e+00 | **** | 0.2880 | small | 27.3 | 62.3 | 2.28 |
| 500km | Not Reef-supporting | Reef-supporting | 1384 | 635 | 96299.0 | 0.00e+00 | **** | 0.3760 | moderate | 38.4 | 89.7 | 2.34 |
| 1000km | Not Reef-supporting | Reef-supporting | 1069 | 569 | 61175.5 | 0.00e+00 | **** | 0.4530 | moderate | 40.7 | 115.0 | 2.83 |
| 2000km | Not Reef-supporting | Reef-supporting | 790 | 518 | 37679.0 | 0.00e+00 | **** | 0.5140 | large | 48.5 | 146.0 | 3.01 |
| **Excl. Unlithified/Poorly-lithified & Sieved** |  |  |  |  |  |  |  |  |  |  |  |  |
| **Excl. Unknown Lithification Style** |  |  |  |  |  |  |  |  |  |  |  |  |
| 100km | Not Reef-supporting | Reef-supporting | 1559 | 476 | 105482.5 | 0.00e+00 | **** | 0.2580 | small | 22.0 | 41.5 | 1.89 |
| 500km | Not Reef-supporting | Reef-supporting | 1021 | 501 | 63157.0 | 0.00e+00 | **** | 0.3560 | moderate | 29.6 | 69.0 | 2.33 |
| 1000km | Not Reef-supporting | Reef-supporting | 818 | 480 | 44273.5 | 0.00e+00 | **** | 0.3980 | moderate | 32.0 | 78.4 | 2.45 |
| 2000km | Not Reef-supporting | Reef-supporting | 613 | 448 | 26495.5 | 0.00e+00 | **** | 0.4810 | moderate | 35.3 | 105.0 | 2.97 |
| **Excl. Unlithified/Poorly-lithified & Sieved** |  |  |  |  |  |  |  |  |  |  |  |  |
| **Incl. Unknown Lithification Style** |  |  |  |  |  |  |  |  |  |  |  |  |
| 100km | Not Reef-supporting | Reef-supporting | 2149 | 585 | 148448.0 | 0.00e+00 | **** | 0.3000 | moderate | 26.5 | 58.7 | 2.22 |
| 500km | Not Reef-supporting | Reef-supporting | 1356 | 633 | 93328.0 | 0.00e+00 | **** | 0.3740 | moderate | 37.5 | 86.8 | 2.31 |
| 1000km | Not Reef-supporting | Reef-supporting | 1046 | 566 | 58392.5 | 0.00e+00 | **** | 0.4490 | moderate | 40.2 | 110.0 | 2.74 |
| 2000km | Not Reef-supporting | Reef-supporting | 780 | 514 | 35102.0 | 0.00e+00 | **** | 0.5220 | large | 47.8 | 143.0 | 2.99 |
| **Lithified Deposits Only** |  |  |  |  |  |  |  |  |  |  |  |  |
| 100km | Not Reef-supporting | Reef-supporting | 1430 | 460 | 94204.0 | 0.00e+00 | **** | 0.2740 | small | 20.9 | 40.3 | 1.93 |
| 500km | Not Reef-supporting | Reef-supporting | 945 | 492 | 57874.5 | 0.00e+00 | **** | 0.3540 | moderate | 28.8 | 59.3 | 2.06 |
| 1000km | Not Reef-supporting | Reef-supporting | 757 | 473 | 39258.5 | 0.00e+00 | **** | 0.4080 | moderate | 30.7 | 73.0 | 2.38 |
| 2000km | Not Reef-supporting | Reef-supporting | 570 | 444 | 24873.0 | 0.00e+00 | **** | 0.4780 | moderate | 34.3 | 95.5 | 2.78 |
| **Non-lithified Deposits Only** |  |  |  |  |  |  |  |  |  |  |  |  |
| 100km | Not Reef-supporting | Reef-supporting | 273 | 59 | 3957.0 | 9.73e-01 | ns | 0.0022 | small | 38.0 | 39.4 | 1.04 |
| 500km | Not Reef-supporting | Reef-supporting | 182 | 87 | 3929.0 | 3.84e-01 | ns | 0.0613 | small | 40.5 | 45.6 | 1.13 |
| 1000km | Not Reef-supporting | Reef-supporting | 159 | 95 | 3128.0 | 4.78e-02 | * | 0.1470 | small | 38.1 | 59.7 | 1.57 |
| 2000km | Not Reef-supporting | Reef-supporting | 114 | 95 | 2294.0 | 9.56e-03 | ** | 0.2070 | small | 39.0 | 64.2 | 1.65 |
| **Unknown Lithification Style Only** |  |  |  |  |  |  |  |  |  |  |  |  |
| 100km | Not Reef-supporting | Reef-supporting | 861 | 275 | 41469.0 | 8.80e-06 | **** | 0.1610 | small | 23.3 | 37.3 | 1.60 |
| 500km | Not Reef-supporting | Reef-supporting | 633 | 375 | 34694.5 | 4.00e-07 | **** | 0.2030 | small | 27.6 | 42.0 | 1.52 |
| 1000km | Not Reef-supporting | Reef-supporting | 517 | 364 | 23875.5 | 0.00e+00 | **** | 0.2600 | small | 28.2 | 49.6 | 1.76 |
| 2000km | Not Reef-supporting | Reef-supporting | 405 | 339 | 13701.0 | 0.00e+00 | **** | 0.3630 | moderate | 33.1 | 78.4 | 2.37 |

**Table S3:**
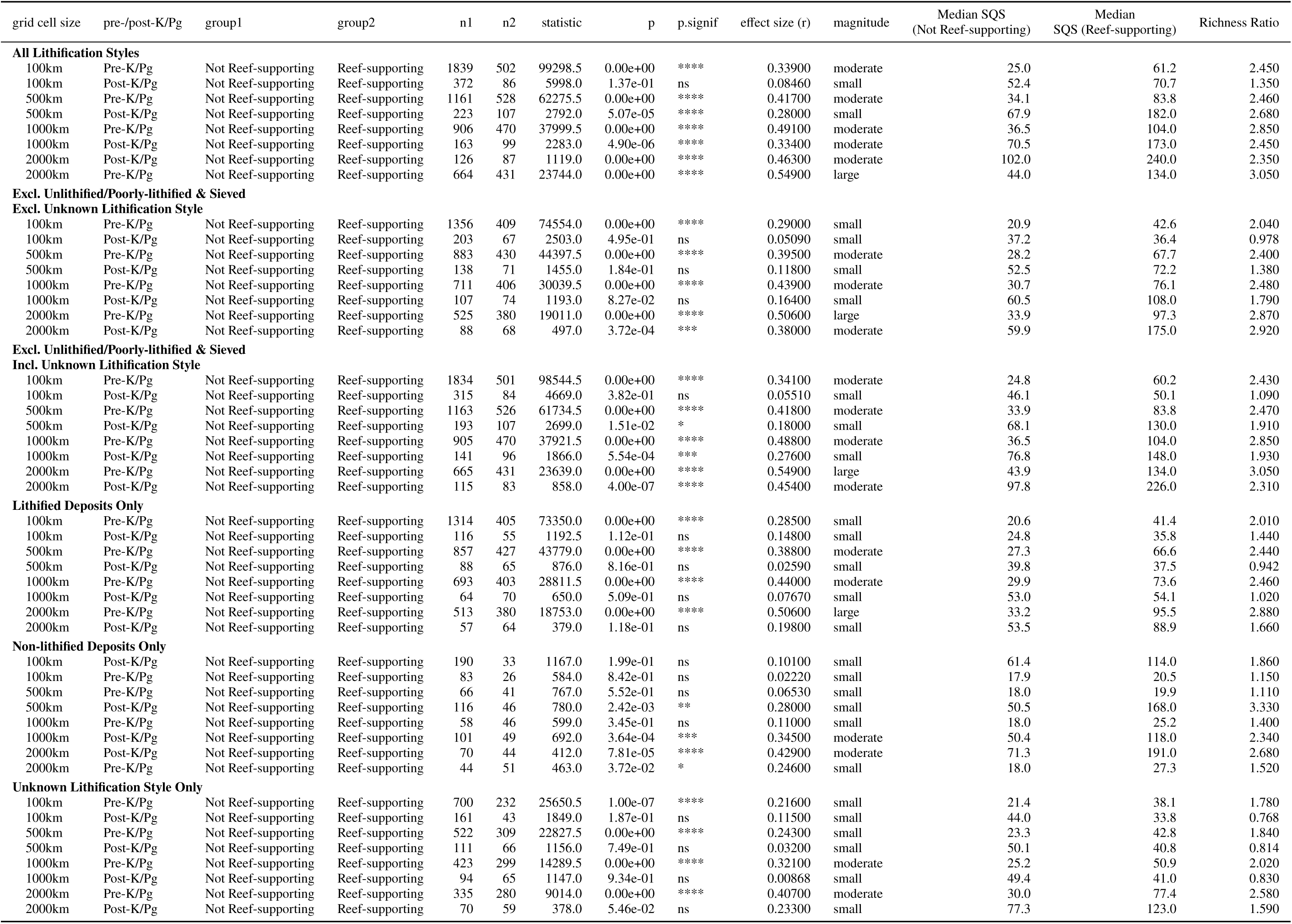
Wilcoxon tests of whether there is a statistically significant difference in diversity among reef-supporting/non-reef-supporting regions, within pre- and post-K/Pg intervals, for several different sifting criteria (see Supplementary Methods) and equal-area hexagonal/pentagonal grid-cell sizes (spacings of 100 km, 500 km, 1000 km and 2000 km). Magnitude denotes strength of effect size.

**Table S4:**
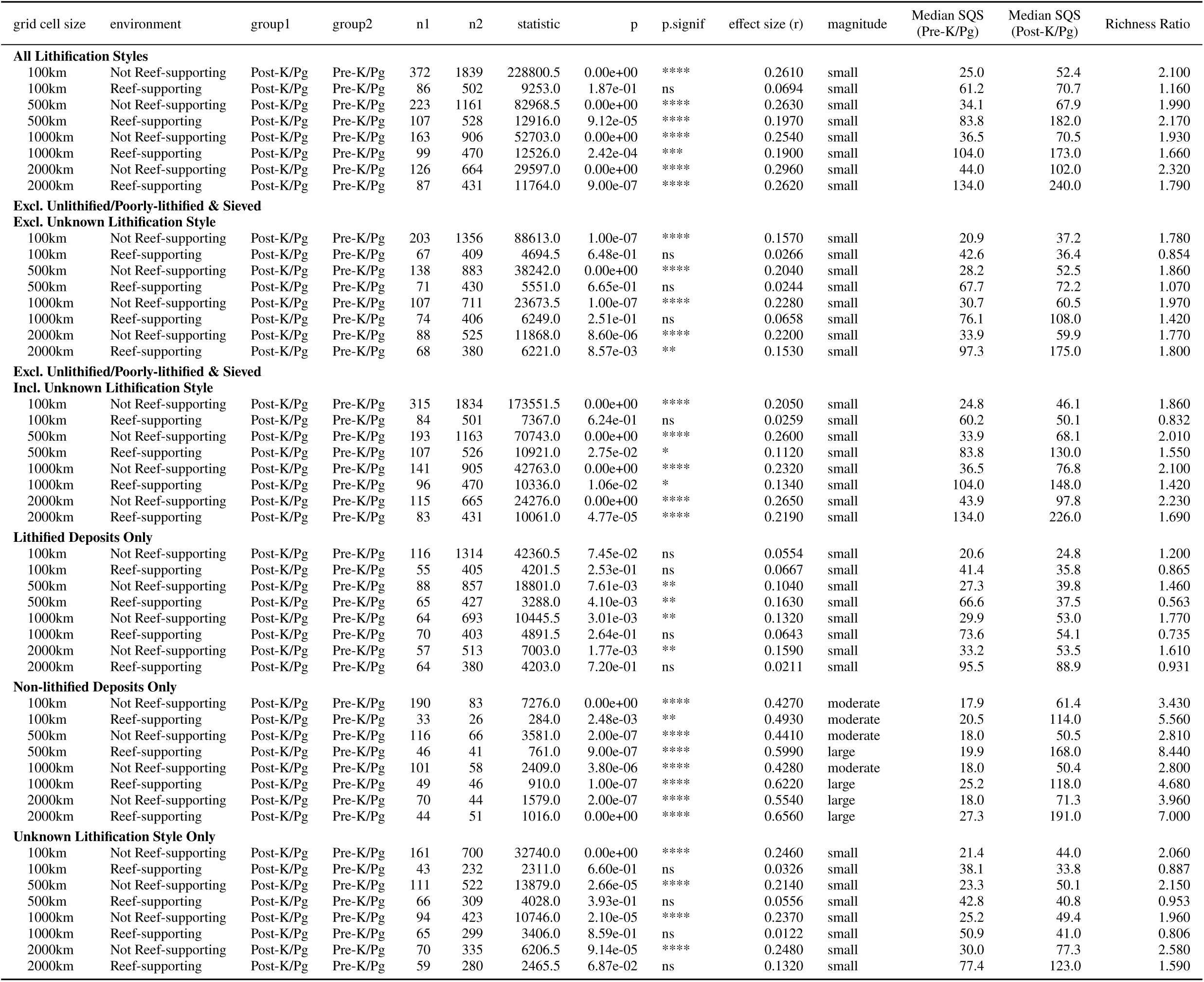
Wilcoxon tests of whether there is a statistically significant difference in diversity between the pre- and post-K/Pg intervals within reef-supporting/non-reef-supporting regions, for several different sifting criteria (see Supplementary Methods) and equal-area hexagonal/pentagonal grid-cell sizes (spacings of 100 km, 500 km, 1000 km and 2000 km). Magnitude denotes strength of effect size.

**Table S5:** Effects of gastropods on K/Pg diversity increase (testing whether diversity is greater in Cz than it is in Pz-Mz)

| environment | group1 | group2 | n1 | n2 | statistic | p | p.signif | effect size (r) | magnitude | Median SQS<br>(Post-K/Pg) | Median SQS<br>(Pre-K/Pg) | Richness Ratio |
| --- | --- | --- | --- | --- | --- | --- | --- | --- | --- | --- | --- | --- |
| **Gastropods** |  |  |  |  |  |  |  |  |  |  |  |  |
| Not Reef-supporting | Pre-K/Pg | Post-K/Pg | 134 | 78 | 1895.0 | 0.00e+00 | **** | 0.5310 | large | 44.0 | 11.5 | 3.83 |
| Reef-supporting | Pre-K/Pg | Post-K/Pg | 123 | 41 | 1152.5 | 2.00e-07 | **** | 0.4060 | moderate | 75.3 | 23.4 | 3.22 |
| **Marine Invertebrates** |  |  |  |  |  |  |  |  |  |  |  |  |
| Not Reef-supporting | Pre-K/Pg | Post-K/Pg | 826 | 134 | 38605.0 | 0.00e+00 | **** | 0.1810 | small | 44.3 | 23.2 | 1.91 |
| Reef-supporting | Pre-K/Pg | Post-K/Pg | 470 | 108 | 24303.5 | 4.92e-01 | ns | 0.0286 | small | 59.6 | 48.8 | 1.22 |
| **Marine Invertebrates**<br>**Excluding Gastropods** |  |  |  |  |  |  |  |  |  |  |  |  |
| Not Reef-supporting | Pre-K/Pg | Post-K/Pg | 812 | 125 | 42301.5 | 2.71e-03 | ** | 0.0980 | small | 34.7 | 22.7 | 1.53 |
| Reef-supporting | Pre-K/Pg | Post-K/Pg | 464 | 104 | 25082.5 | 5.28e-01 | ns | 0.0265 | small | 50.8 | 47.2 | 1.08 |

**Table S6:**
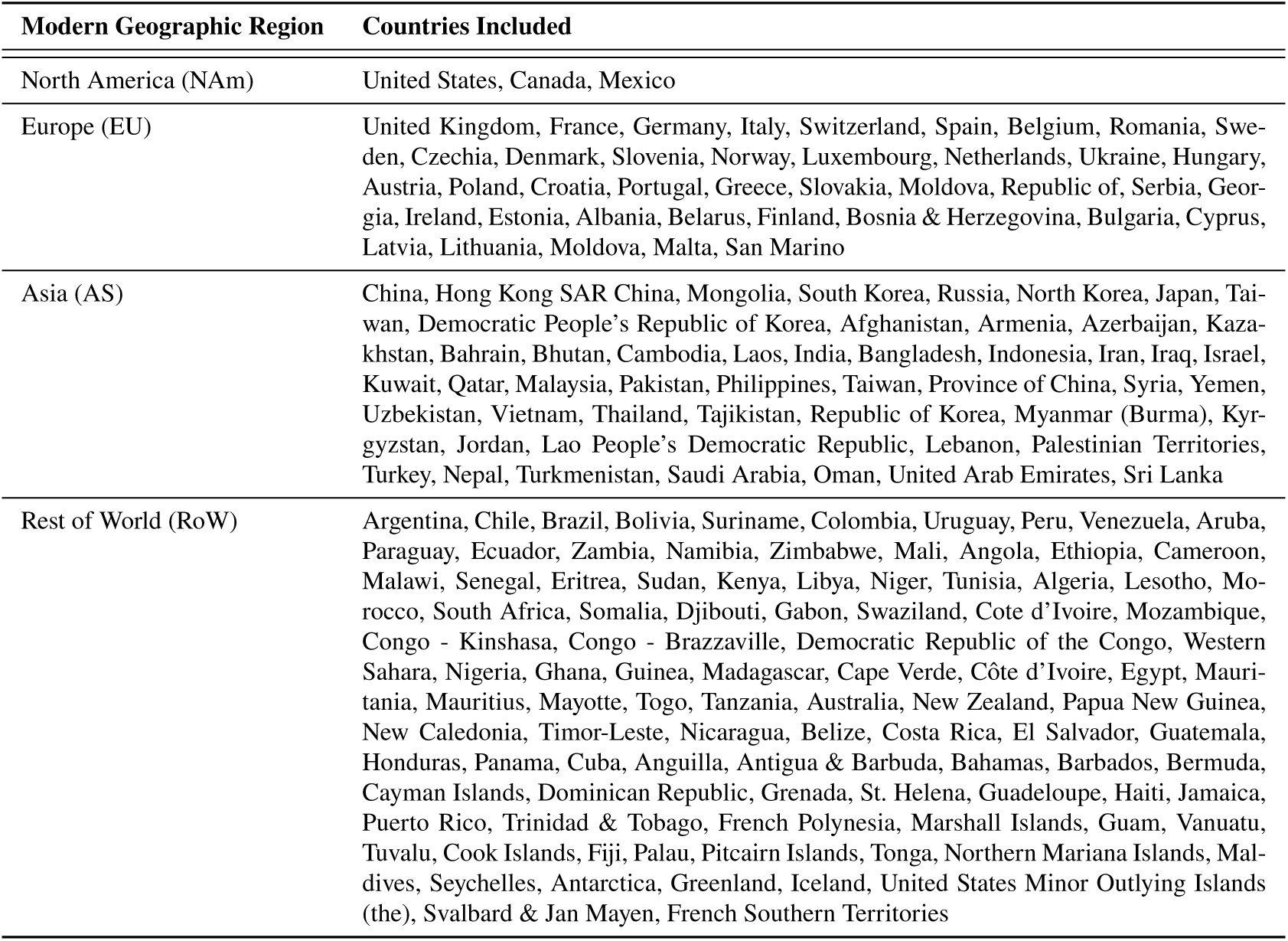
Definitions of modern geographic regions.

**Table S7:**
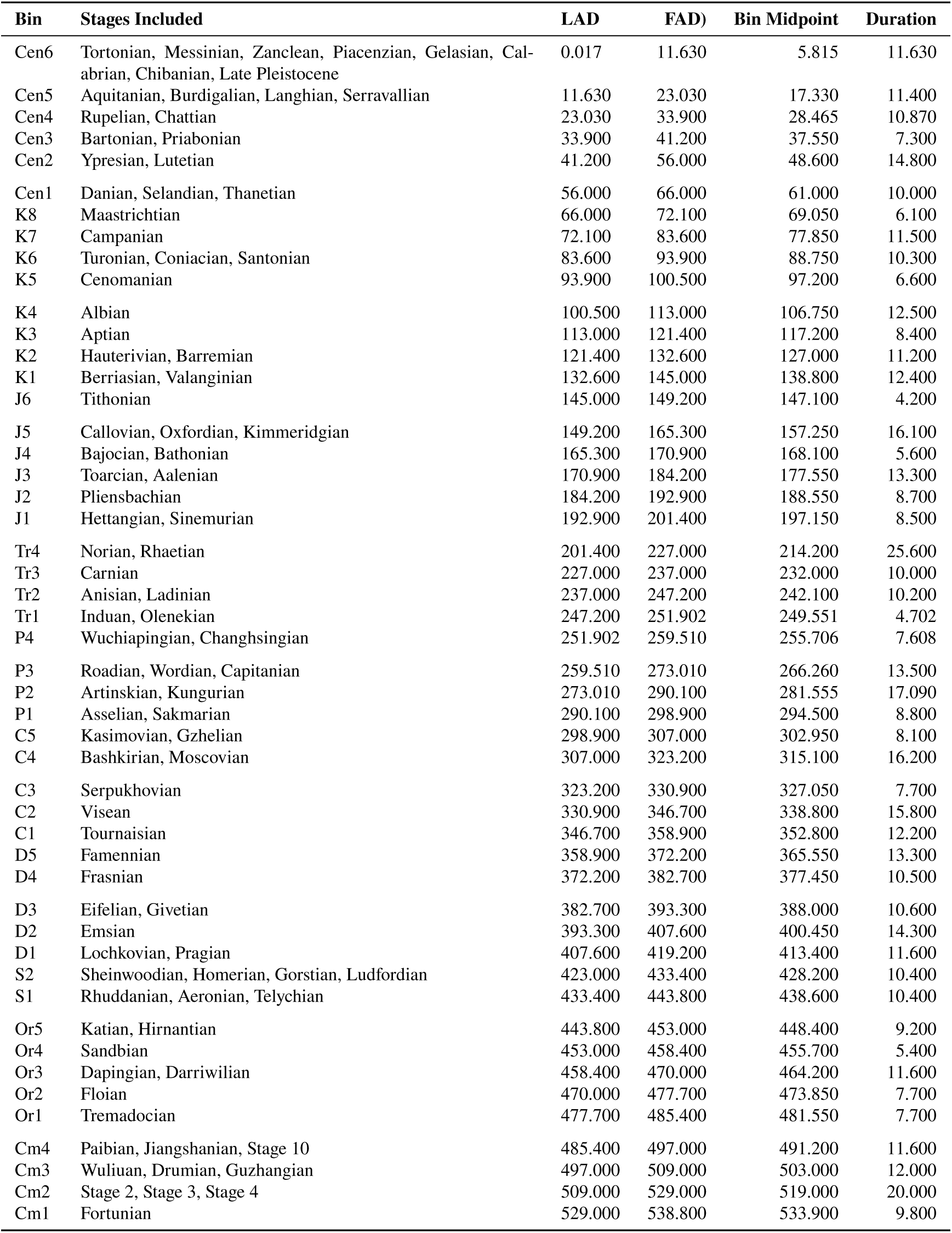
Definitions of equal-length time bins.

## References and Notes

1. M. L. Reaka-Kudla, The Global Biodiversity of Coral Reefs: A Comparison with Rain Forests, Biodiversity II (Joseph Henry Press), pp. 83–108 (1997).

2. N. Knowlton, et al., Coral Reef Biodiversity (John Wiley & Sons, Ltd), chap. 4, pp. 65–78 (2010), 10.1002/9781444325508.ch4, https://onlinelibrary.wiley.com/doi/abs/10.1002/9781444325508.ch4.

3. R. Fisher, et al., Species Richness on Coral Reefs and the Pursuit of Convergent Global Estimates. Current Biology 25 (4), 500–505 (2015), doi:10.1016/j.cub.2014.12.022.

4. R. K. Bambach, Species Richness in Marine Benthic Habitats through the Phanerozoic. Paleobiology 3, 152–167 (1977).

5. A. M. Bush, R. K. Bambach, Did Alpha Diversity Increase during the Phanerozoic? Lifting the Veils of Taphonomic, Latitudinal, and Environmental Biases. The Journal of Geology 112 (6), 625–642 (2004), doi:10.1086/424576.

6. D. Jablonski, K. Roy, J. W. Valentine, Out of the Tropics: Evolutionary Dynamics of the Latitudinal Diversity Gradient. Science 314 (5796), 102–106 (2006), doi:10.1126/science.1130880.

7. W. Kiessling, C. Simpson, M. Foote, Reefs as Cradles of Evolution and Sources of Biodiversity in the Phanerozoic. Science 327 (5962), 196–198 (2010), doi:10.1126/science.1182241.

8. J. Alroy, Geographical, Environmental and Intrinsic Biotic Controls on Phanerozoic Marine Diversification. Palaeontology 53 (6), 1211–1235 (2010), doi:10.1111/j.1475-4983.2010.01011.x.

9. V. J. Roden, et al., Fossil Liberation: A Model to Explain High Biodiversity in the Triassic Cassian Formation. Palaeontology 63 (1), 85–102 (2020), doi:10.1111/pala.12441.

10. R. A. Close, R. B. J. Benson, E. E. Saupe, M. E. Clapham, R. J. Butler, The Spatial Structure of Phanerozoic Marine Animal Diversity. Science 368 (6489), 420–424 (2020), doi:10.1126/science.aay8309.

11. R. Wood, The Changing Biology of Reef-Building. PALAIOS 10 (6), 517 (1995), doi:10.2307/3515091.

12. W. Kiessling, Long-Term Relationships between Ecological Stability and Biodiversity in Phanerozoic Reefs. Nature 433 (7024), 410–413 (2005), doi:10.1038/nature03152.

13. W. Renema, et al., Hopping Hotspots: Global Shifts in Marine Biodiversity. Science 321 (5889), 654–657 (2008), doi:10.1126/science.1155674.

14. J. J. Sepkoski, A Factor Analytic Description of the Phanerozoic Marine Fossil Record. Paleobiology 7 (1), 36–53 (1981), doi:10.1017/s0094837300003778.

15. J. Alroy, The Shifting Balance of Diversity among Major Marine Animal Groups. Science 329 (5996), 1191–1194 (2010), doi:10.1126/science.1189910.

16. J. Alroy, et al., Phanerozoic Trends in the Global Diversity of Marine Invertebrates. Science 321 (5885), 97–100 (2008), doi:10.1126/science.1156963.

17. R. B. Benson, R. Butler, R. A. Close, E. Saupe, D. L. Rabosky, Biodiversity across Space and Time in the Fossil Record. Current Biology 31 (19), R1225–R1236 (2021), doi:10.1016/j.cub.2021.07.071.

18. The Paleobiology Database (http://www.paleobiodb.org) .

19. For Further Details, See Supplementary Information.

20. A. Chao, L. Jost, Coverage-based Rarefaction and Extrapolation: Standardizing Samples by Completeness Rather than Size. Ecology 93 (12), 2533–2547 (2012), doi:10.1890/11-1952.1.

21. M. G. Powell, M. Kowalewski, Increase in Evenness and Sampled Alpha Diversity through the Phanerozoic: Comparison of Early Paleozoic and Cenozoic Marine Fossil Assemblages. Geology 30 (4), 331 (2002), doi: 10.1130/0091-7613(2002)030⟨0331:iieasa⟩2.0.co;2.

22. J. J. Sepkoski, A Kinetic Model of Phanerozoic Taxonomic Diversity. III. Post-Paleozoic Families and Mass Extinctions. Paleobiology 10 (02), 246–267 (1984), doi:10.1017/s0094837300008186.

23. A. Rojas, J. Calatayud, M. Kowalewski, M. Neuman, M. Rosvall, A Multiscale View of the Phanerozoic Fossil Record Reveals the Three Major Biotic Transitions. Communications Biology 4 (1), 309 (2021), doi:10.1038/s42003-021-01805-y.

24. P. Bouchet, P. Lozouet, P. Maestrati, V. Heros, Assessing the Magnitude of Species Richness in Tropical Marine Environments: Exceptionally High Numbers of Molluscs at a New Caledonia Site: MOLLUSCAN SPECIES RICHNESS IN A TROPICAL MARINE ENVIRONMENT. Biological Journal of the Linnean Society 75 (4), 421–436 (2002), doi:10.1046/j.1095-8312.2002.00052.x.

25. M. Kowalewski, W. Kiessling, M. Aberhan, F. T. Fürsich, Ecological, Taxonomic, and Taphonomic Components of the Post-Paleozoic Increase in Sample-Level Species Diversity of Marine Benthos. Paleobiology 32 (04), 533–561 (2006), doi:10.1666/05074.1.

26. A. M. Bush, R. K. Bambach, Sustained Mesozoic–Cenozoic Diversification of Marine Metazoa: A Consistent Signal from the Fossil Record. Geology p. G37162.1 (2015), doi:10.1130/g37162.1.

27. W. Kiessling, Geologic and Biologic Controls on the Evolution of Reefs. Annual Review Of Ecology Evolution And Systematics 40 (1), 173–192 (2009), doi:10.1146/annurev.ecolsys.110308.120251.

28. R. Wood, The Ecological Evolution of Reefs. Annual Review of Ecology and Systematics 29 (1), 179–206 (1998), doi:10.1146/annurev.ecolsys.29.1.179.

29. T. P. Hughes, et al., Coral Reefs in the Anthropocene. Nature 546 (7656), 82–90 (2017), doi:10.1038/nature22901.

30. T. P. Hughes, et al., Global Warming and Recurrent Mass Bleaching of Corals. Nature 543 (7645), 373–377 (2017), doi:10.1038/nature21707.

31. O. Hoegh-Guldberg, Climate Change, Coral Bleaching and the Future of the World’s Coral Reefs. Marine and Freshwater Research (1999), doi:10.1071/MF99078.

32. O. Hoegh-Guldberg, et al., Coral Reefs Under Rapid Climate Change and Ocean Acidification. Science 318 (5857), 1737–1742 (2007), doi:10.1126/science.1152509.

33. J. Alroy, Effects of Habitat Disturbance on Tropical Forest Biodiversity. Proceedings of the National Academy of Sciences 16, 201611855–16 (2017), doi:10.1073/pnas.1611855114.

34. C. R. Scotese, An Atlas of Phanerozoic Paleogeographic Maps: The Seas Come In and the Seas Go Out. Annual Review of Earth and Planetary Sciences 49 (1), 1–50 (2021), doi:10.1146/annurev-earth-081320-064052.

35. W. H. Berger, F. L. Parker, Diversity of Planktonic Foraminifera in Deep-Sea Sediments. Science 168 (3937), 1345–1347 (1970), doi:10.1126/science.168.3937.1345.

36. W. Kiessling, M. Krause, PARED – An online database of Phanerozoic reefs (2022), https://www.paleo-reefs.pal.uni-erlangen.de.

37. A. J. W. Hendy, The Influence of Lithification on Cenozoic Marine Biodiversity Trends. Paleobiology 35 (01), 51–62 (2009), doi:10.1666/07047.1.

38. A. J. W. Hendy, Taphonomy, Process and Bias Through Time. Topics in Geobiology pp. 19–77 (2010), doi: 10.1007/978-90-481-8643-3 2.

39. A. D. Hawkins, M. Kowalewski, S. Xiao, Breaking down the Lithification Bias: The Effect of Preferential Sampling of Larger Specimens on the Estimate of Species Richness, Evenness, and Average Specimen Size. Paleobiology 44 (02), 326–345 (2018), doi:10.1017/pab.2017.39.

40. G. Daley, A. Bush, The Effects of Lithification on Fossil Assemblage Biodiversity and Composition: An Experimental Test. Palaeontologia Electronica (2020), doi:10.26879/1119.

41. R. Nawrot, Decomposing Lithification Bias: Preservation of Local Diversity Structure in Recently Cemented Storm-Beach Carbonate Sands, San Salvador Island, Bahamas. PALAIOS 27 (3), 190–205 (2012), doi:10.2110/palo.2011.p11-028r.

42. J. Alroy, et al., Effects of Sampling Standardization on Estimates of Phanerozoic Marine Diversification. Proceedings of the National Academy of Sciences 98 (11), 6261–6266 (2001), doi:10.1073/pnas.111144698.

43. Ádám T. Kocsis, N. B. Raja, chronosphere: Evolving Earth System Variables (2023).

44. R. Barnes, R-Barnes/dggridR.

45. R. A. Close, S. W. Evers, J. Alroy, R. J. Butler, How Should We Estimate Diversity in the Fossil Record? Testing Richness Estimators Using Sampling-Standardised Discovery Curves. Methods in Ecology and Evolution 28, 1023–15 (2018), doi:10.1111/2041-210x.12987.

46. T. C. Hsieh, K. H. Ma, A. Chao, iNEXT: An R Package for Rarefaction and Extrapolation of Species Diversity (Hill Numbers). Methods in Ecology and Evolution pp. 1–6 (2016), doi:10.1111/2041-210x.12613.

47. I. J. Good, The Population Frequencies of Species and the Estimation of Population Parameters. Biometrika 40 (3-4), 237–264 (1953), doi:10.1093/biomet/40.3-4.237.

48. R. A. Close, et al., Diversity Dynamics of Phanerozoic Terrestrial Tetrapods at the Local-Community Scale. Nature Ecology & Evolution 3 (4), 590–597 (2019), doi:10.1038/s41559-019-0811-8.

49. J. Cohen, Statistical Power Analysis. Current Directions In Psychological Science 1 (3), 98–101 (1992).

